# Streptolysin O Accelerates the Conversion of Plasminogen to Plasmin

**DOI:** 10.1101/2024.05.15.594458

**Authors:** Di Tang, Hamed Khakzad, Elisabeth Hjortswang, Lars Malmström, Simon Ekström, Lotta Happonen, Johan Malmström

## Abstract

*Streptococcus pyogenes* is a human-specific bacterial pathogen that produces several proteins that exploit the plasminogen (PLG)-plasmin (PLM) fibrinolysis system to dismantle blood clots and to facilitate spread and survival within the host. In this study, we explored the interactions between streptolysin O (SLO), a key cytolytic toxin produced by *S. pyogenes*, and human plasma proteins using affinity-enrichment mass spectrometry. SLO binds specifically to PLG, and this interaction accelerates the conversion of PLG to PLM by tissue-type plasminogen activator (tPA). To further investigate the molecular detail of the PLG-SLO interaction, we employed hydrogen/deuterium exchange mass spectrometry combined with targeted cross-linking mass spectrometry, uncovering that SLO binding induces local conformational shifts in PLG. These changes lead to the formation of a stabilized intermediate PLG-SLO complex that becomes significantly more sensitive to proteolytic processing by tPA. Our findings reveal a conserved moonlighting pathomechanistic role for SLO extending beyond the well characterized cytolytic activity. Additionally, the work underscores the diversity of functional proteomics in identifying and clarifying new host-pathogen interactions.

## Introduction

*Streptococcus pyogenes*, or Group A streptococcus (GAS), is a gram positive human bacterial pathogen that can cause a wide range of infections. Worldwide, GAS is responsible for 767 million superficial infections such as pyoderma and pharyngitis, 1.78 million severe infections, and over 500,000 deaths annually.^1^ Infections caused by GAS can additionally lead to autoimmune-associated sequalae such as rheumatic fever/heart disease and poststreptococcal acute glomerulonephritis.^2^ Recently, several reports have observed antibiotics resistant clinical GAS isolates^3,4^, which represents a substantial public health concern due to the high prevalence of GAS infections. These observations emphasize the need for more in-depth understanding of GAS pathogenesis to promote the development of innovative therapeutic strategies.

Infections caused by GAS activates the coagulation system leading to the formation of a fibrin network to entrap the bacteria.^5^ As a consequence, GAS has developed the capacity to exploit the human haemostatic system to promote microbial survival and dissemination. In particular, several mechanisms to control the fibrinolysis pathway and the plasminogen (PLG)/plasmin (PLM) system have evolved in GAS to induce fibrinolysis and to escape entrapment.^6^ PLG circulates as two different glycosylated proteoforms (type I: Asn_308_, Thr_365_; type II: Thr_365_).^7^ The mature form of PLG is a 90 kDa protein, subdivided into 7 consecutive domains (**Figure 1A**). The first domain is referred to as the Pan Apple N-terminal (PAN) domain, followed by five homologous Kringle domains (K1-K5), and a peptidase S1 domain (PSD). Circulating PLG adopts a tight conformation held together by intramolecular inter-domain interactions to prevent aberrant activation under normal conditions.^8^ Upon binding to fibrin or cellular receptors, a competitive binding occurs with the lysine-binding sites (LBS) on the PLG, leading to a conformational change in PLG to expose its activation loop.^9^ This activation loop is subsequently cleaved by plasminogen activators (PAs) to form active plasmin^10^, where the two cleaved chains remain interconnected by disulfide bonds. Cleavage of the activation loop and proteolytic removal of the PAN domain leads to the formation of mature plasmin. Small-angle X-ray scattering (SAXS) techniques have revealed the conformational transition of PLG from its closed to open form, showing a single-step conversion process that results in multiple final conformations with high flexibility in solution.^9^ However, interfering signals from macromolecular PLG-binding partners, such as bacterial proteins and cellular receptors, makes SAXS ineffective in detecting these changes in more complex conditions. Currently, several PAs have been characterized, including tissue-type plasminogen activator (tPA), urokinase-type plasminogen activator (uPA), kallikrein, and factor XII (Hageman factor)^11^, as well as several bacterial proteins from different species.^12^

**Figure 1.**
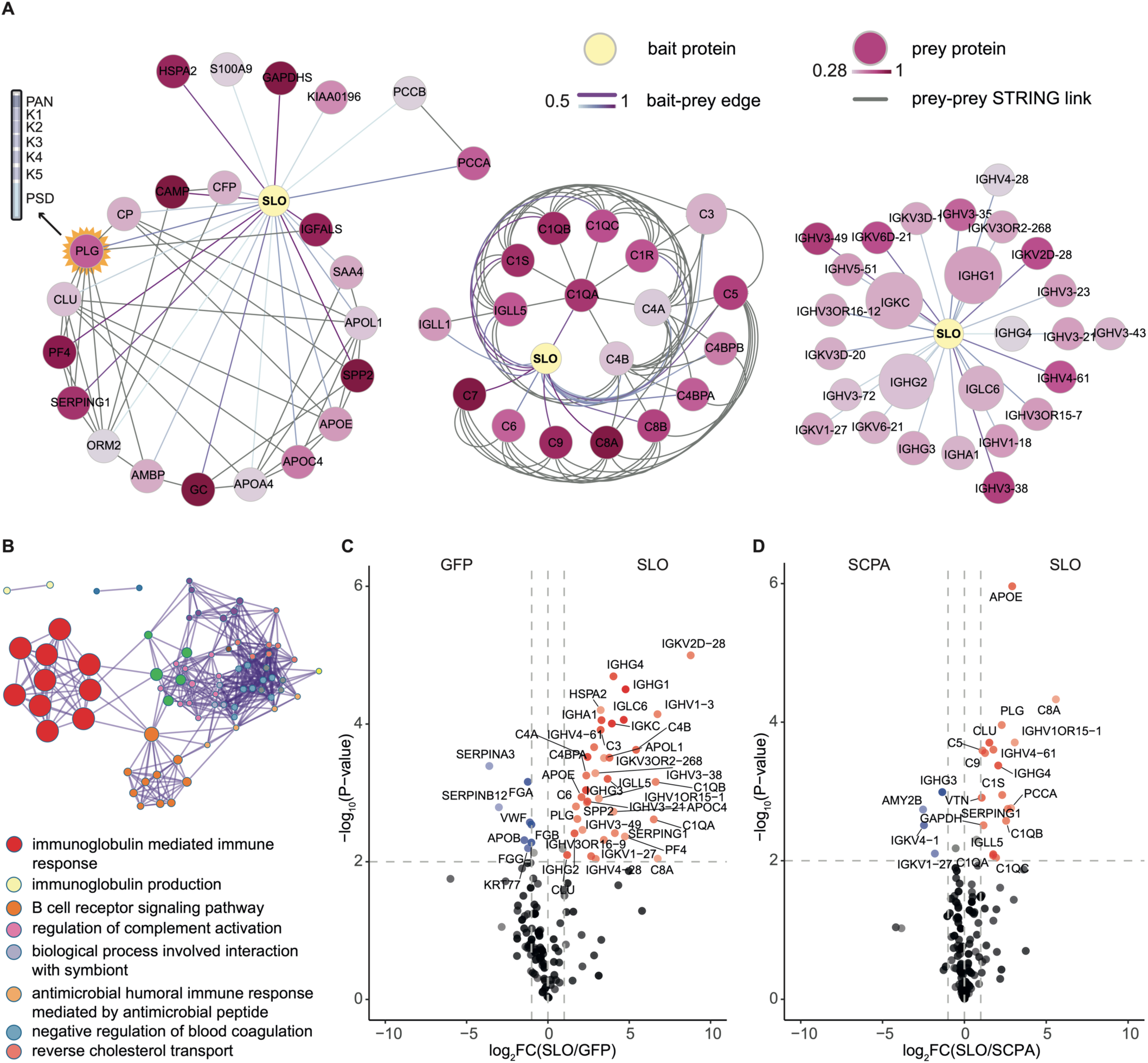
A protein-protein interaction network between SLO and human plasma proteins. Recombinantly tagged SLO (streptolysin O) and SCPA (C5a peptidase) baits were used to enrich proteins from undiluted pooled human plasma in three independent replicates, followed by data independent acquisition mass spectrometry (DIA-MS) analysis. The affinity enriched (AE)-DIA quantification data was sorted into a bait-prey matrix for scoring by MiST and visualisation in Cytoscape. **A)** This panel of networks depict the interactions between SLO and plasma proteins. Nodes represent proteins, with size and colour indicating abundance and specificity of the enriched prey protein, while connecting edges, represent protein-protein interactions (PPI). The width and colour of the edges are based on MiST scores. The SLO-plasma PPI network further integrates high-confidence PPI data from the STRING interactome database, identifying associations among prey proteins and categorizing them into subnetworks: left, network of other plasma proteins; middle, network of complement components; right: network of immunoglobulins, **B)** A network of enriched Biological Process (GO BP) terms, derived from 68 prey proteins, organized into clusters. The most representative term from each of the top 8 clusters were used as annotations. Volcano plots illustrate statistical protein abundance comparisons between **C)** SLO and GFP, and **D)** SLO and SCPA, with fold changes plotted against -log_10_ P values from multiple t-tests on extracted ion intensities. Dashed lines define the significance threshold of |fold change| >1 and an adjusted p-value < 0.01. Proteins surpassing this threshold in the SLO comparison are marked with red dots, those in the GFP/SCPA comparison in blue, and non-significant isolated proteins in black. All significantly enriched proteins are labelled with gene names.

So far, six streptococcal proteins have been reported to interact with PLG, such as PLG-binding GAS M-like proteins (PAM)^13^, α-enolase (SEN)^14^, gyceraldhehyde3-phosphate-dehydrogenase (GAPDH)^15^, streptococcal surface dehydrogenase (SDH)^16^, extracellular protein factor (Epf)^17^ and streptokinase (SKA) ^18^. Streptokinase is a well-defined PLG-binding streptococcal protein that forms a 1:1 activator complex with PLG to catalyse other free PLG molecules to active Plasmin^19^. PAM binds to PLG between the K1-K2 domains using the a1a2 motif to introduce a relaxed conformation that renders PLG more susceptible to activation. Notably, a similar enhanced conversion has been observed in the binding of α-Enolase to PLG.^20^ The PAM-mediated increase in PLM production can occur either through SKA- cooperated activation or directly by exploiting host plasminogen activators (PAs).^21^ Importantly, previous reports have shown that GAS virulence was enhanced in a human-PLG- transgene mouse model^22^, demonstrating that the interaction between GAS and PLG is critically important for GAS pathogenicity, dissemination, and survival.^23^

Streptolysin O (SLO) is a 60 kDa pore-forming toxin secreted by nearly all GAS strains. The first 69 N-terminal residues of SLO forms a disordered region, followed by three discontinuous domains (D1-D3) and a membrane-binding domain (D4) at the C terminus.^24^ SLO primarily binds to cholesterol-rich membranes and oligomerizes to form a pre-pore followed by D3- driven transition into a pore to cause cytolysis.^25,26^ Beyond this main biological function, SLO also acts as an immune-modulatory molecule for neutrophils^27^ and impairs phagocytic clearance of GAS.^28^ Recent studies on other members of the cholesterol-dependent cytolysin (CDC) family proteins such as pneumolysin (PLY)^29^ and perfringolysin O (PFO)^30^ have shown that this protein family has several biological functions beyond pore-formation, like reducing inflammation or binding to glycans. Other studies also indicate that there are other non-classic cholesterol CDC cellular receptors and CDC-binding plasma proteins like human albumin.^31,32^ SLO is present in more than 98% of the sequenced GAS genomes with a relatively low sequence variability of less than 2%^33^, making it a suitable target for developing therapeutic interventions.

In this study, we applied affinity-enrichment mass spectrometry (AE-MS) to investigate the interaction network formed between SLO and human plasma proteins, revealing that SLO binds specifically to PLG. In contrast to Streptokinase, the binding of SLO to PLG did not result in direct activation of PLG but rendered PLG more sensitive to tPA activation. Using an integrative structural mass spectrometry strategy combining hydrogen/deuterium mass spectrometry (HDX-MS), targeted cross-linking mass spectrometry (TX-MS)^34^ and structural modelling, we demonstrated that SLO binding to PLG leads to a stabilized intermediate complex, which facilitates tPA in catalysing increased plasmin production to benefit the course of GAS infection.

## Results

### A protein-protein interaction network between SLO and human plasma proteins

We have previously shown that many virulence factors produced by GAS form protein networks with human plasma and saliva proteins.^35^ To further expand these interaction networks, we produced two virulence factors from GAS in recombinant form, streptolysin O (SLO) and C5a peptidase (SCPA), genetically fused with an affinity tag. These tagged bait proteins were used to affinity-enrich (AE) interacting human proteins from pooled healthy plasma, which were identified by high-resolution LC-MS/MS (AE-MS) using tagged green fluorescence protein (GFP) as a reference control. The two GAS antigens enriched distinct set of human plasma proteins as seen in the heatmap and clustering analysis in the **Extended Data Figure 1A-B**. The proteins enriched by the respective tagged proteins were organized into a bait-prey matrix and processed using the MiST workflow.^36,37^ The resulting network was visualized using Cytoscape^37,38^ and demonstrated 90 plasma proteins interacting with the two GAS bait proteins, forming 116 bait-prey edges. Notably, 42 of these interactions were exclusively formed with SLO (**Extended Data Figure 2A**). Integrating the SLO-centric network with the high-confidence interactions from the STRING database^39^ expanded the network to a total of 203 edges, revealing subnetworks predominantly comprised of plasma proteins, complement components, and immunoglobulins (**Figure 1A**).

A prominent feature of the interaction networks was the enrichment of several immunoglobulin G (IgG) chains that were specifically enriched by SLO or SCPA along with components of the complement system that are known to interact with the fragment crystallizable (Fc) part of IgG, such as C1q. This observation shows that the pooled healthy human plasma contains multiple IgG clones specific for either SLO or SCPA, which was corroborated using enzyme-linked immunosorbent assay (ELISA) (**Extended Data Figure 2B**). Functional over-representation analysis of the SLO enriched plasma proteins revealed a network of terms such as immunoglobulin associated functions, regulation of the complement activation and negative regulation of blood coagulation (**Figure 1B**, **Extended Data Figure 3A-B**). In the next step, we used FDR-adjusted multiple t-test comparing SLO/GFP and SLO/SCPA to identify significant protein-protein interactions specific for SLO using a threshold of >1.0-fold change and an adjusted p-value of <0.01 (**Figure 1C-D**). The differential analyses consistently identified apolipoprotein E (APOE), plasminogen (PLG), clusterin (CLU), vitronectin (VTN) and plasma protease C1 inhibitor (SERPING1) to be the most significant interactions that were specific for SLO. It is noteworthy that these interactions were formed even in the presence of anti-SLO IgGs, suggesting that SLO-specific IgG does not outcompete these interactions.

To further filter out weak binders to SLO and to imitate the microenvironment formed during plasma leakage, the pooled human plasma was diluted, and the AE-MS experiments were repeated twice. The putative SLO-specific interactors identified above were quantitatively monitored in the pulldown samples from all three baits (SLO, SCPA and GFP) across three plasma dilutions. In these experiments, we observe that SLO enriches certain IgG chains and complement components significantly more than GFP and SCPA under all dilution conditions. Consistent across the undiluted and the 50% diluted conditions, PLG, APOE, and CLU were significantly enriched by SLO when compared to SCPA and GFP separately, whereas only PLG was significantly enriched in the 10% diluted condition (**Figure 2A-B**, **Extended Data Figure 4A-B**). In conclusion, we used AE-MS to map two plasma protein interaction networks formed around SLO or SCPA and uncovered three binding partners that specifically interacted with SLO.

**Figure 2.**
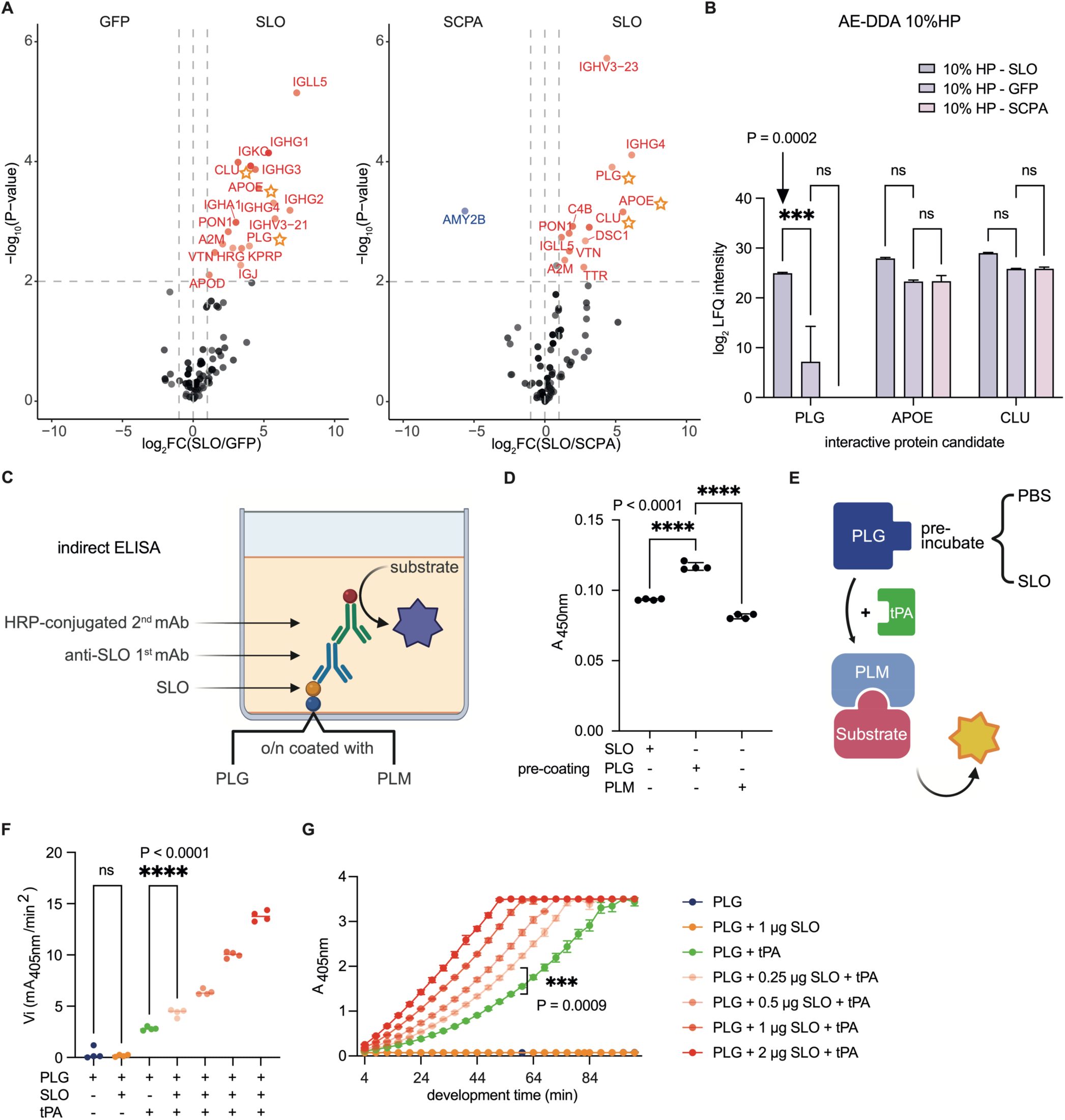
SLO binds specifically to PLG and enhances tPA-mediated conversion of plasminogen to plasmin. The pooled human plasma was diluted and analysed through AE-DDA workflow in three independent replicates to identify proteins that interact specifically to streptolysin O compared to the GFP/SCPA control background. **A)** Differentially SLO-enriched protein was determined by normalized label-free quantification (LFQ) using fold-change and multiple t-test as cut-offs. The same significance threshold and colour scheme as Figure 1 were applied here. The consistently enriched proteins were PLG (Plasminogen), APOE (apolipoprotein E) and CLU (clusterin) and are marked with stars. **B)** Two-way ANOVA group analysis on log2-transformed LFQ intensities of the three SLO-interacting plasma protein candidates. **C-D)** An indirect ELISA was used to quantify binding of SLO to PLG/PLM (plasmin), with higher absorbance values indicating stronger binding. **E)** Schematic outline of the plasminogen activation assay. The assay involves pre-incubating PLG with/without SLO, followed by the addition of tissue-type plasminogen activator (tPA) and a chromogenic PLM substrate. The absorbance was continuously measured and reflects the amount of new plasmin that was generated correlating to PLG activation rate by tPA. **F)** An initial PLG activation velocity (Vi) comparison was made using one-way ANOVA test across all conditions. “+”: with; “- “: without. **G)** Connected dot plot shows the cumulative effect of PLG activation over time and tested for statistical significance using a two-way ANOVA test. Significance levels are indicated as “ns” (no significance), “*”, “**”, “***”, and “****”, corresponding to p-values of < 0.1, < 0.01, < 0.001, and < 0.0001, respectively.

### SLO binds specifically to PLG and enhances tPA-mediated conversion of plasminogen to plasmin

Zymogen plasminogen circulates under normal conditions in a tight conformation resistant to proteolytic activation to prevent the formation of PLM.^40^ However, upon binding to fibrin or cell surfaces, PLG shifts to a more relaxed conformation, enabling rapid conversion to active plasmin (PLM) by one of the plasminogen activators.^10^ To first determine if SLO binds to both PLG and PLM, we employed an indirect ELISA assay as PLG and PLM have the same primary structure and are indistinguishable by mass spectrometry analysis. The wells were coated with equal amount of either PLG or PLM, followed by incubation with SLO. The level of SLO binding was then quantified using an anti-SLO monoclonal antibody and an HRP-conjugated secondary antibody (**Figure 2C**). The results revealed that SLO binds only to PLG (**Figure 2D**). To determine if SLO can bind to both of main isoforms of human Plasminogen^7^ (type I: Asn_308_, Thr_365_; type II: Thr_365_), we employed targeted glycoproteomics analysis to search glycopeptides in the AE-MS DDA datasets. Glycopeptides containing N-linked Asn_308_ or O-linked Thr_365_ were confidently identified in all SLO-enriched plasma pulldowns, indicating that SLO can enrich both predominant proteoforms of PLG (**Supplementary Data Figure 1A**).

In contrast to Streptokinase, our initial experiments showed that SLO does not directly catalyse PLG to PLM. Consequently, we next investigated if binding of SLO could alter PLG sensitivity to tPA activation in a similar fashion as fibrin or cell surfaces. In these experiments, PLG was incubated with SLO followed by the addition of tPA and a chromogenic PLM substrate (**Figure 2E**). The initial activation rates were calculated and shown as individual dot points for four replicates across seven conditions (**Figure 2F**). Additionally, kinetic mode analysis was conducted to measure the accumulation of the end-product (chromophore para-nitroaniline) over a 120 min incubation period, until the positive control (PLG + tPA) reached saturation (**Figure 2G**). The results show that SLO significantly enhanced the tPA-mediated conversion in a dose-dependent manner without affecting PLM activity (**Supplementary Data Figure 1B**). Furthermore, we determined that the binding of PLG does not affect the haemolytic function of SLO, suggesting that these two interactions can co-occur and are not mutually exclusive (**Supplementary Data Figure 1C**).

### HDX-MS reveals protected regions and protein dynamics of PLG upon SLO binding

To map the binding interfaces between PLG and SLO and to investigate the dynamic aspects of the interaction, we performed HDX-MS to explore changes in deuterium uptake in PLG upon binding to SLO. In HDX-MS, increased or decreased exchange of deuterium uptake measured by mass spectrometry indicates interaction interfaces or changes in protein state/conformation during for example protein-protein binding or protein-small molecule drug interactions.^41^ PLG was analysed in both the apo (unbound) and the SLO-bound states, where HDX-MS identified 150 common peptides of high-confidence annotation, achieving 65.3% sequence coverage with many overlapping peptides (**Extended Data Figure 5A**). Changes in deuterium uptake (ΔDU) across five deuteration intervals (0, 30, 300, 3000, and 9000 s) for each peptide were summed and visualized in a butterfly plot (**Figure 3A**). Overall, the ΔDU between apo PLG and SLO-bound PLG were marginal, with notable exceptions in the peptide regions spanning residues 74-84 in PAN, 385-438 in K4 domain, and 716-737 and 768-793 in PSD. PLG peptides with significantly decreased deuterium uptake determined by FDR-controlled significance test, also pinpointed several protected regions in K1, K2, and K3 upon SLO binding, indicating a global protein stabilization during the longest deuteration interval (**Figure 3B**). Shorter deuterium labelling times (<=300 s) are more likely to reveal changes associated with initial binding.^42^ During these labelling times, regions 34-48 and 80-88 of PAN, 405-421 in K4, and 726-734 in PSD were protected indicating that these regions are likely involved in the initial binding (**Extended Data Figure 5B**). The more rigorous hybrid significance test further highlighted the most significantly protected regions (192-203 and 277- 288 in K2, 658-669 in PSD), as shown in the volcano plot comparing the highest deuteration interval of 9000 s (**Fig. 3C)**. Representative overlapping protected peptides across all five deuteration intervals demonstrating the temporal protection are shown in **Figure 3D** and **Extended Data Figure 5C**. To provide additional insights into PAN and PSD domains, the average differential deuterium uptake change at the residue level for these regions were plotted in a barcode plot (**Figure 3E**). Notably, the region spanning around 650-670 in PSD transitioned to a more protected conformation over time possibly due to an allosteric effect while the protection of the proximate regions 720-730 and 795-810 in PSD decreased with time. In contrast, the regions 620-630 and 770-790 in PSD were consistently protected across all labelling intervals. Decreased protection was observed for regions 35-50 and 75-90 in PAN. A full barcode plots depicting both apo and SLO-bound state mean/differential deuterium uptake is shown in the **Extended Data Figure 5D**. The HDX-MS data corroborated the direct interaction between SLO and plasminogen protein, and pinpointed protected regions on the PLG molecule due to SLO-binding or allosteric effect. Importantly, the absence of confident deprotected peptides also suggests that PLG does not undergo major relaxation or conformational rearrangement when bound to SLO. However, based on HDX data, we could not confidently differentiate local SLO-binding interfaces from global domain stabilization.

**Figure 3.**
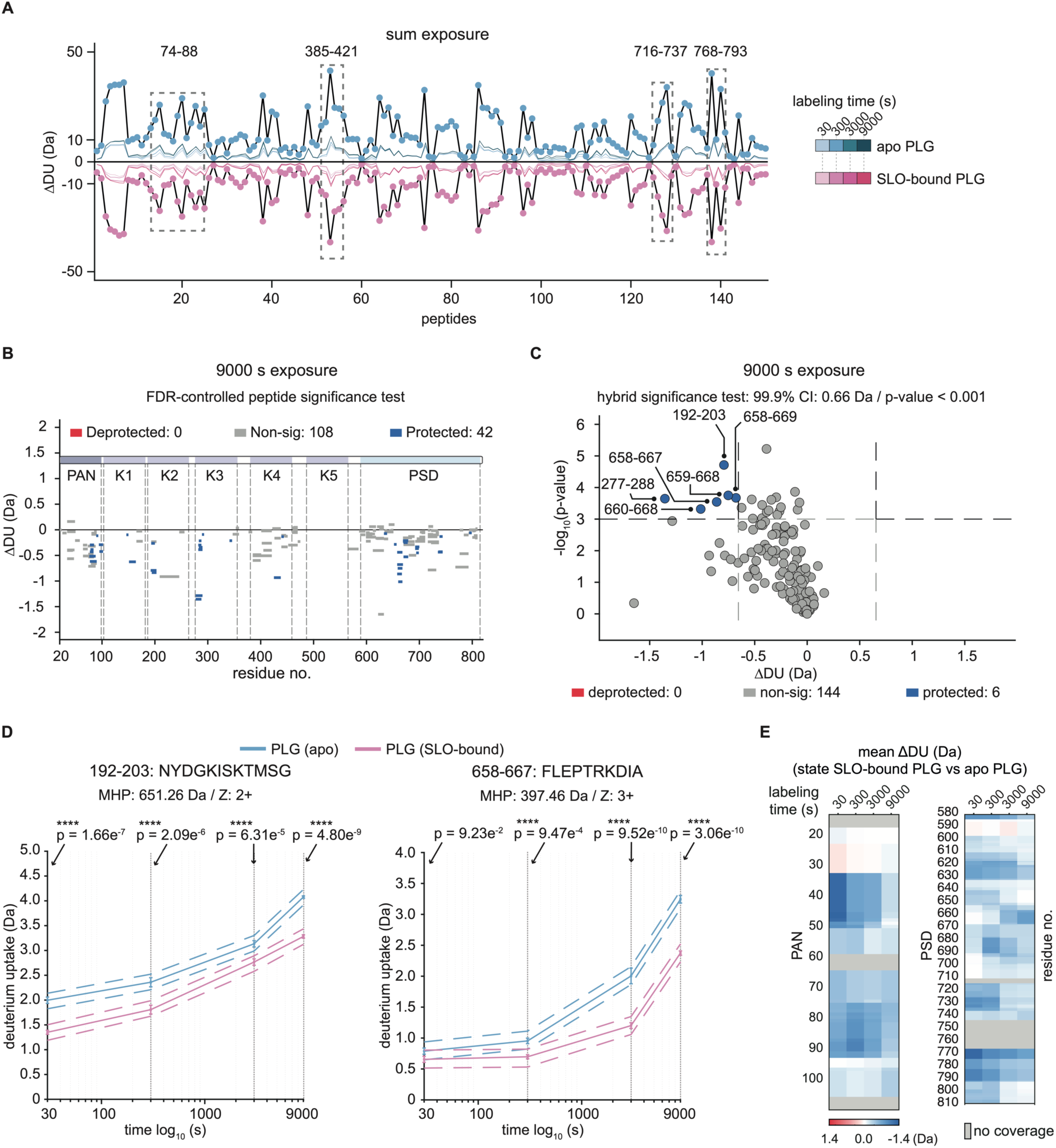
HDX-MS reveals protected regions and protein dynamics of PLG upon SLO binding. **A)** The butterfly plot aggregates the summed deuterium uptake of all 150 identified PLG peptides over the four labelling intervals of two states (apo and SLO-bound), with each shared peptide represented by a dot. A dashed-line box highlights protected regions of PLG upon SLO binding, annotated with residue numbers. **B)** A Woods plot illustrates the deuterium uptake change of each PLG peptide after 9000 s deuteriation, where the length of each line corresponds to peptide length, mapped across the PLG sequence. FDR-corrected significance test (confidence level of 99.9%) was applied and indicates protected peptides in blue, deprotected in red, and non-significant changes in grey, correlated to the corresponding PLG domains. **C)** A volcano plot showing protected and deprotected peptide after 9000 s deuteriation, with a threshold of > 0.66 Da change and p-values < 0.001. Each peptide is depicted as a dot and coloured. **D)** Kinetic plots for two representative protected peptides as 192-203 and 658-667, with significance testing across four labelling times, color-coded by the different states (apo and SLO-bound). Significance levels “ns” for non-significant and “****” for >99.9% significance are marked, with p-values. ’MHP’ refers to theoretical molecular weight of the peptide, Z for charge state. **E)** The barcode plot illustrates the mean change in deuterium uptake for PAN and PSD domain residues when comparing apo (unbound) and SLO-bound states. Each residue is represented with a colour gradient indicating the extent of change as a function of labelling interval.

### XL-MS defines interaction sites and protein dynamics of the PLG-SLO interaction

We speculated that the increased sensitivity to tPA activation could possibly be linked to local conformation shifts in PLG molecule upon SLO binding. To further elucidate the binding interfaces within PLG-SLO complex and to assess the protein dynamics in both binding partners, we performed in-solution cross-linking mass spectrometry (XL-MS). Both disuccinimidyl suberate (DSS) and disuccinimidyl glutarate (DSG) were used to cross-link either PLG or PLM to SLO. In addition, DSS was used to cross-link PLG with tPA, or PLG together with tPA and SLO to investigate possible transient heterodimeric or heterotrimeric complexes. DSG has a spacer arm length of 7.7 Å compared to DSS of 11.4 Å, which generates shorter distance constraints between the cross-linkable reactive lysine residues in proximity. The cross-linked peptides were searched primarily using pLink2^43^, and MaxLynX.^44^ From four independent PLG-SLO cross-linked datasets, we identified in total 53 DSS and 78 DSG inter-protein cross-linked spectrum matches (CSMs) with pLink2, and 75 DSS and 69 DSG inter-protein CSMs with MaxLynX at an FDR of 0.01 (**Supplementary Data Table 1**). Linkage maps were created using XlinkCyNET based on the pLink2-searched results. These linkage maps highlight the 16 DSS and 22 DSG cross-linked sites formed between PLG and SLO (**Figure 4A-B**) and that only one cross-linked site was identified between PLM and SLO supported by only one CSM. These results corroborate the ELISA result and prove that SLO does not bind strongly to PLM. In addition, we identified 5 inter-links between tPA and PLG and 12 inter-links between tPA, PLG and SLO supporting the existence of intermediate state during tPA catalysation (**Extended Data Figure 6A**). Most of the inter-protein cross-linked sites were identified in the three primary Pfam-annotated domains in PLG (Lys81 in PAN, Lys433 in K4 domain, Lys664 and Lys727 in PSD) (**Extended Data Figure 6B-C**). Reciprocally for SLO, most inter-links were identified in domains D1, D3, and D4. Representative MS/MS spectra of DSS or DSG XL peptide pairs between PLG and SLO are shown in the **Extended Data Figure 6D-F, G-I**. Additionally, DSS inter-links were identified between tPA and PLG (**Extended Data Figure 6J**), as well as between tPA and SLO (**Extended Data** Figure 6K) when cross linking was performed on all three proteins. Inter- links can be formed between tPA and PLG due to the slow kinetics of tPA-mediated plasminogen activation in solution, whereas the identification of tPA-SLO inter-links indicates the formation of intermediate heterotrimeric complex for enhanced catalysation on PLG.

**Figure 4.**
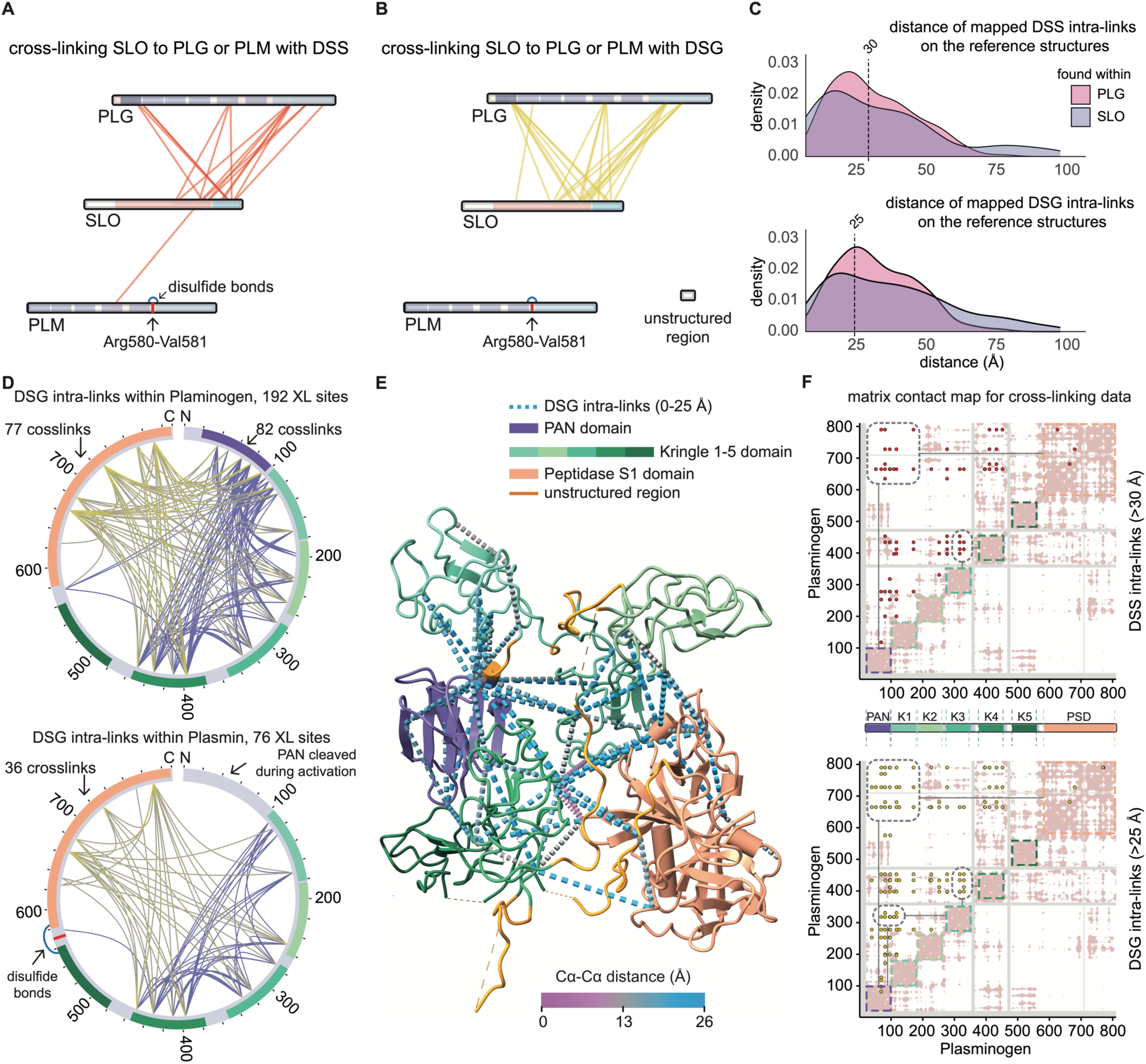
XL-MS defined interaction sites and protein dynamics of the PLG-SLO interaction. Linkage maps were constructed using Cytoscape with all identified inter-protein cross-links that passed the 1% false discovery rate (FDR) threshold. The cross-linked sites between SLO and PLG/PLM, with **A)** DSS inter-links in red and **B)** DSG inter-links in yellow, and protein family domains coloured in segments accordingly. Cleavage site for conversion of PLG to PLM is depicted as a red segment, as well as disulfide bonds connecting two chains after the cleavage. **C)** Overlapping density plots display the Cα-Cα distance measurement distribution of all mapped DSS and DSG intra-protein cross-links found within PLG or SLO. **D)** Two circular plots show the DSG intra-protein cross-linked sites as lines within PLG or PLM protein, with cross-links connecting the peptidase domain highlighted in yellow shading. **E)** A crystal structure (PDB: 4DUR) of PLG, coloured by domain, with DSG intra-links (< 25 Å) found within SLO-bound PLG protein shown in dot-line style pseudo bonds. **F)** Two matrix contact maps showing the distance-violating intra-protein cross-links (> 30 Å for DSS; > 25 Å for DSG) as dots for PLG. White background represents the amino acid sequence of PLG on the both the x- and y-axis. Grey colour indicates absence of residue on the structure and pink for two residues in close proximity (Cα-Cα distance within 0-25 Å).

To investigate changes in the protein conformation/dynamics in PLG and SLO during binding, we examined in the next step the pLink2-reported intra-links within PLG, PLM, and SLO. These non-overlapping intra-links were mapped onto the reference structures (PLG, PDB: 4DUR; SLO, PDB: 4HSC) previously solved by X-ray crystallography. The distances between the backbone Cα atoms of the cross-linked lysine pairs were then measured and corresponding CSM counts summarized in bar plots and circos plots found in the **Extended Data** Fig.7A**-D**. Displaying the density plots of the distances between the Cα atoms for SLO and PLG separately, demonstrates that fraction of the intra-links with overlength Cα- Cα distance was considerably higher in SLO compared to the fraction of overlength Cα-Cα distance found within PLG (**Figure 4C**). As there have been no reports regarding major conformational changes or rearrangement of SLO in its solution phase, these results rather indicate that SLO is more dynamic in nature or can adopt a higher degree of domain movement when bound to PLG.

When comparing the number of intra-links in PLG to those within plasmin (PLM), we observed a notable reduction of intra-links found within PLM, from 192 to 76 cross-linked sites, and from 1177 to 322 CSM counts, most likely due to relaxation of the molecule after tPA-cleavage (**Figure 4D**). The loss of intra-links, which only form when neighbouring domains are in close proximity, suggests particularly greater mobility of PSD domain within PLM compared to PLG. Mapping the PLG intra-links within 0-25 Å distance measurement to the reference PDB crystal structure, together with HDX data, suggests that the PLG conformation remains largely unchanged without any major structural rearrangement upon SLO binding (**Figure 4E**). However, 75 DSS intra-links and 129 DSG intra-links still violated the allowed maximum distance in the context of PLG crystal structure. Plotting these overlength cross-links in a matrix contact map revealed that they primarily clustered in the PAN, PSD and K4 domains (**Figure 4F**). This pattern indicates that there are movement in these domains when PLG interacts with SLO. A similar mapping and matrix contact map on SLO also suggests that in particular domain 4 was subjected to increased movement when SLO binds to PLG (**Extended Data Figure 8A-D**). Taken together, these findings indicate that PLG undergoes local conformational changes mainly driven by PAN, K4 and PSD during its interaction with SLO. Furthermore, XL-MS perfectly overlapped with the protected regions (75-90 in PAN; 650-670, 720-730 and 770-790 in PSD) in PLG identified by HDX.

### TX-MS predicts PLG-SLO pairwise model and illustrates the binding interfaces

In the final step, targeted cross-linking mass spectrometry (TX-MS)^34^ was performed using all the distance constraints to determine the most important residues involved in the binding interfaces between PLG and SLO. TX-MS builds pairwise models of the protein complex using the Rosetta docking protocol^45^ and considers how well each model fulfils the inter-links between the two interacting proteins. Initial tests using AlphaFold2-Multimer^46^ without integrating the XL- or HDX-derived distance constraints resulted in a protein complex shown in **Figure 5A**. According to this model, the domain 4 of SLO binds to peptidase S1 domain of PLG via its membrane binding motif. This binding interface aligns with the binding site determined by both HDX-MS and TX-MS. However, the AlphaFold model could not predict the correct orientation of SLO with respect to PLG or explain the inter-links detected between PLG and other SLO regions like domain 3. To clarify this, we first investigated the intramolecular crosslinks of SLO as shown in **Extended Data** Figure 7. Interestingly, we observed multiple intra-links that were inconsistent with both the monomeric and the dimeric form of SLO, suggesting that SLO undergo a conformational change. Computational modelling of these changes was conducted using normal mode-based geometric simulations by NMSim^47^. This analysis indicated that the domain 4 of SLO can move inward forming a bend conformation upon binding to PLG that could explain most of the over length intra-links (K145-K540, K189-K407, K491-K540, K154-K417, K196-K298, K407-K540, and K407-K491) (**Figure 5B**).

**Figure 5.**
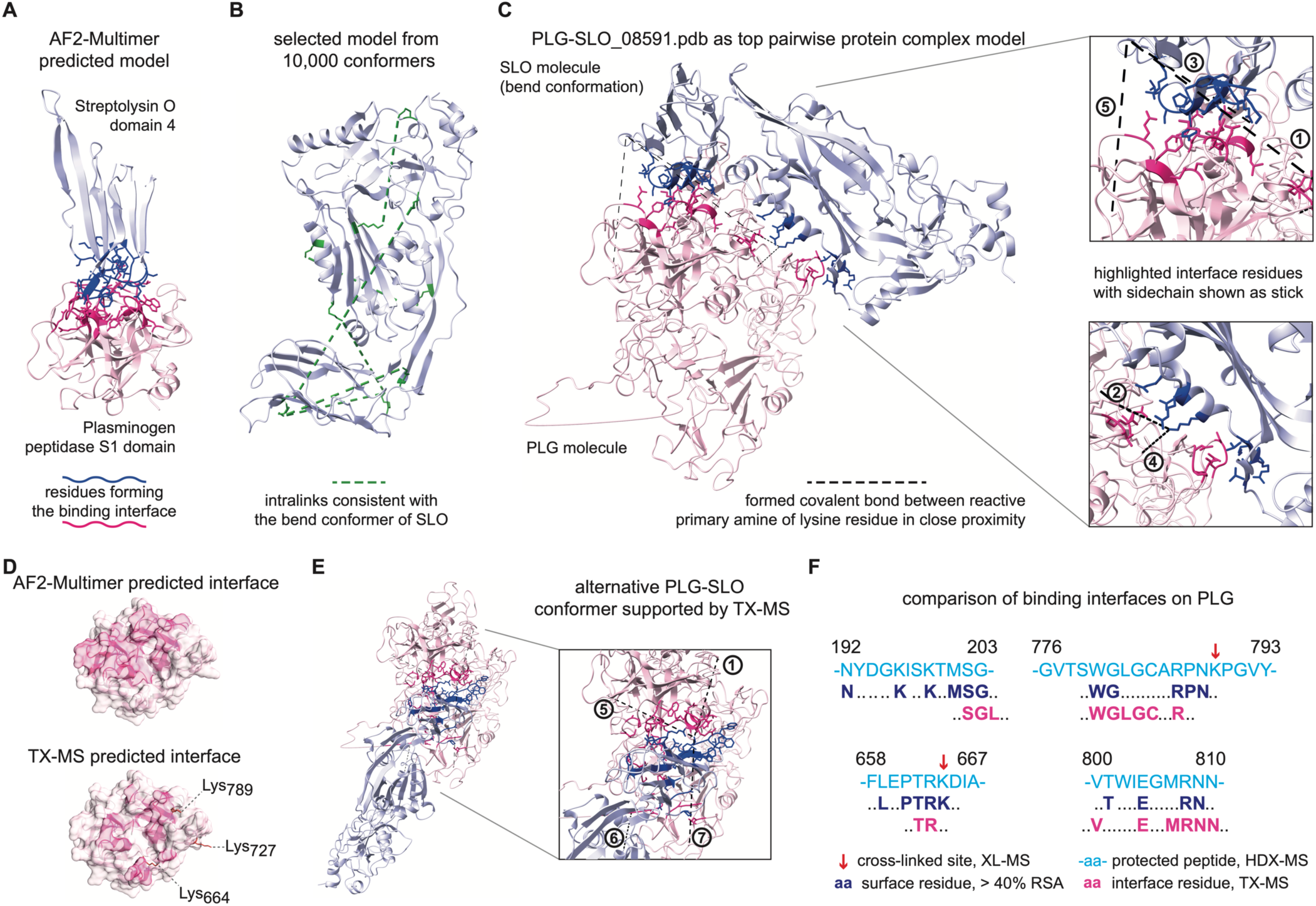
TX-MS predicts PLG-SLO pairwise model and illustrates the binding interfaces. **A)** An AlphaFold-Multimer predicted pairwise complex model of the SLO domain 4 and the plasminogen PSD domain. The binding interfaces were determined by PRODIGY, highlighted in different colours respectively and shown in stick representation for the residue side chains. **B)** The bend conformer of SLO protein was modelled by normal mode simulations, accommodating intra-links which were inconsistent with the SLO crystal structure (PDB: 4HSC). **C)** The top 1 pairwise model as PLG-SLO_08591 is presented, with the two binding interfaces highlighted in darker colours. The upper one between the PLG PSD and SLO domain 4 is supported by four cross-links, while the lower one between PLG PSD & K2 and SLO domain 3 is supported by 2 cross-links. Covalent DSS linker bonds are visualized as dashed lines, connecting the reactive lysine side chains. **D)** A PLG PSD binding interface comparison shown in surface representation between AlphaFold2-Multimer predicted and TX-MS models. **E)** An alternative set of cross-links suggests another possible pairwise PLG-SLO model supported by 4 inter-links. **F)** Comparison between contacting interface residues of PLG-SLO complex and four representative HDX-MS derived protected peptides. The Interface residues derived from TX-MS models align with the contact residues proposed by HDX-MS filtered by a relative solvent accessibility (RSA) value of > 40%. Cross-linked sites marked by red arrow .

Next, in order to provide molecular insights into the PLG-SLO complex, two sets of docking models were made using either the static crystal structure of SLO (PDB: 4HSC) or the predicted bend conformer of SLO in **Figure 5B** together with the reference PLG structure (PDB: 4DUR). Here, we selected the model consistent with highest number of intermolecular cross-links with the best docking energy score to build the 1:1 complex (**Figure 5C, Supplementary Data Figure 1D)**. Residues in the binding interface were determined by PRODIGY^48^, highlighted with sidechain in stick presentation and zoomed in with neighbouring covalent bonds formed by linker connecting reactive lysine side chains (**Supplementary Data Table 2**). Intriguingly, the binding interface derived from TX-MS aligned well with the interface predicted by AlphaFold2-Multimer (**Figure 5D**). Still, this top-ranked model was unable to explain all inter- links. Consequently, we cannot exclude that alternative conformers of the PLG-SLO complex may exist (**Figure 5E, Supplementary Data Table 2**), or if two SLO molecules can bind simultaneously to PLG or sequentially at different times.

The HDX-MS results described above identified the binding interfaces at a peptide level resolution. By calculating the relative solvent accessibility (RSA) of each residue in the native PLG structure, likely contact residues were selected based on an RSA of >40%. For the top-ranked model, 6 out of 7 TX-MS predicted interface motifs corroborated the calculated surface-accessible contact residues generated from the HDX-results (**Figure 5F**). Interestingly, without supporting inter-links, the protected peptide 194-203 (also shown in **Figure 3D**) within Kringle 2 domain of PLG was identified by TX-MS. In summary, TX-MS revealed that SLO adopts a bent conformation and enabled the identification of the most important residues involved in the protein binding interface between PLG-SLO. We conclude that leveraging both HDX-MS and XL-MS generates independent evidence that increases the confidence of the identified residues involved in the binding interfaces.

### Highly conserved PLG-binding motifs reveal a moonlighting pathomechanism of SLO

We have previously shown that SLO is produced by 98% of the sequenced GAS genomes with a relatively low sequence variability.^49^ Focusing this analysis on the interfaces required for PLG binding demonstrates that the PLG-binding motifs on SLO are close to 100% conserved (**Extended Data Figure 8E**). These results suggest that the capacity to exploit plasminogen-plasmin system in a streptokinase-independent fashion is conserved in most GAS strains.^50^ Further interrogation with low-frequency mutation and polymorphism within the binding motifs is required to determine the impact of these mutations on the interaction between SLO and PLG.

## Discussion

Here, we demonstrate that streptolysin O (SLO), a well-characterized GAS-secreted pore-forming toxin, binds and alters the conformation of PLG. This interaction makes PLG more sensitive to proteolytic processing by tissue plasminogen activator (tPA) to generate active plasmin (PLM) (**Figure 6A**). Our results show that SLO forms a network of protein interactions with human plasma proteins involved in biological functions related to immune responses and regulation of blood coagulation. Multimodal mass spectrometry and functional analysis further revealed that SLO binds to PLG, leading to increased susceptibility to tPA-mediated formation of PLM. The enhanced tPA-mediated proteolytic conversion was associated with the formation of a more stabilized intermediate state with induced local conformational shifts in the PLG PAN, K4 and PSD domains. These conformational shifts provide increased access for host tPA that accelerates the formation of PLM, supported by the formation of a heterotrimeric PLG- SLO-tPA complex. The tPA-mediated cleavage on PLG further introduced major conformational changes in domain PSD leading to detachment of SLO from the newly generated PLM. Free plasmin then acts to degrade fibrin and dissolve blood clots, while SLO potentially reinitiates the cycle by targeting other inactivated PLG molecules (**Figure 6B**). *In vivo*, we postulate that SLO functionally resembles the co-factor/enhancer role previously shown for PAM (**Figure 6C**), contributing to the SKA-independent PLM generating pathways.

**Figure 6.**
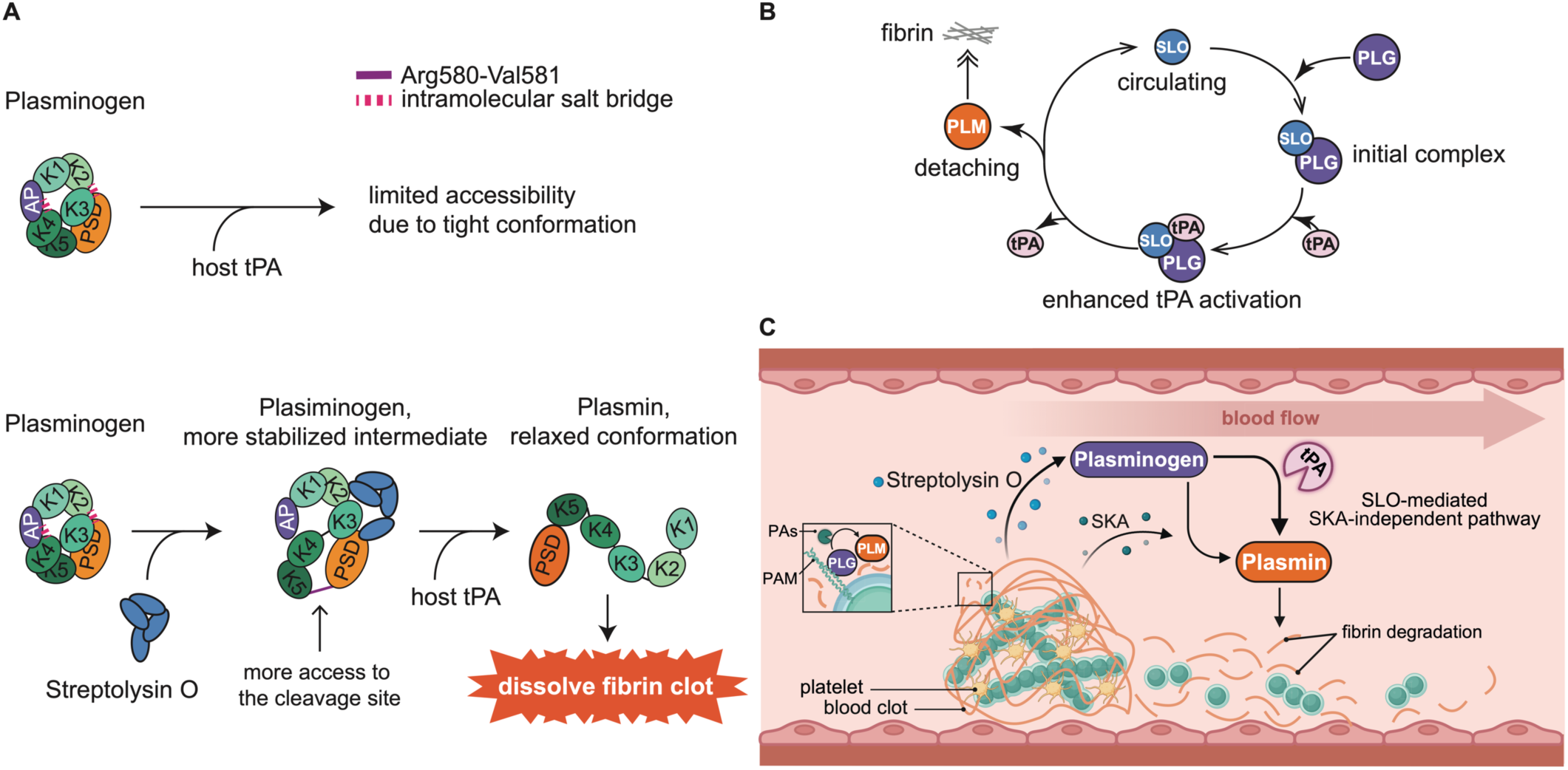
Schematic of SLO-PLG interaction and its implication on *S. pyogenes* pathogenesis. **A)** The plasminogen molecule, depicted in a tight conformation, resists activation to prevent unwarranted enzymatic activity in solution phase. Interaction with GAS-secreted toxin SLO, presumably triggered local conformation shift and stabilizes plasminogen structure. This PLG-SLO intermediate complex then becomes more susceptible to activation by tissue plasminogen activator (tPA), resulting in more plasmin produced. The increased plasmin activity leads to the degradation of the fibrin and extracellular matrix, facilitating blood clot breakdown. **B)** This pathogenic mechanism allows *S. pyogenes* to exploit the host’s fibrinolysis system from distance, promoting deeper tissue invasion and systemic dissemination by utilizing the acquired plasmin activity through the enhanced SLO-mediated activation on Plasminogen. **C)** We hereby propose an SLO-mediated, SKA-independent pathway for tPA activation on human Plasminogen. To evade human host protective fibrin entrapment, *S. pyogenes* has evolved to i) recruit plasminogen and plasmin to its surface, ii) secrete streptokinase to directly catalyse plasmin generation, or iii) acquire increased plasmin activity from distance through the secretion of streptolysin O, thereby enhancing tPA’s conversion of plasminogen to plasmin. PAs: plasminogen activators; PAM: plasminogen-binding group A streptococcal M-like protein; SKA: streptokinase.

Human plasminogen (PLG) plays a central role in the coagulation-fibrinolysis system and is a pathogenic target for several PLG-binding bacterial proteins.^23^ These interactions can modify the activity of PLG to promote bacterial survival by breaking down host fibrin to escape entrapment. To date, six streptococcal proteins have been reported to be Plasminogen-binding, among which streptokinase (SKA) and Plasminogen-binding M-like proteins (PAM) are most studied, which has resulted in detailed characterization of protein-protein binding interfaces.^13,18,19,21,51^ In contrast to SKA, SLO is abundantly produced by in principle all GAS strains. Furthermore, the high degree of conservation in the primary sequence of the SLO binding motifs is in contrast to the high variability observed in the a1a2 motif found in Plasminogen-binding M-like proteins (PAM). Therefore, SLO-mediated enhancement in fibrinolysis transcends serotypes and demonstrates a more conserved and prevalent pathomechanism.

In this study, we used two orthogonal MS-based methods to identify and validate the binding interfaces. The distance constraints generated by XL-MS was further used to guide the downstream modelling of protein complexes. Intriguingly, XL-MS also identified crosslinks between SLO and tPA, suggesting that SLO retains PLG and tPA to catalyse the generation of plasmin (PLM).^11^ In addition, the intra-links identified by XL-MS provided insights into the protein dynamics of both PLG and SLO during interaction. The overlength crosslinks indicate that domain 4 in SLO is subjected to dynamic movement. Similar movement of this domain has previously been reported for SLO-NADase complex.^52^ Given that a crystal structure may not fully represent the dynamic nature of protein movement in solution, the reported crosslinks are likely derived from an ensemble of different conformations, including both bound and unbound states. Reciprocally, the overlength cross-links within PLG indicate local movement in particularly PAN, K4 and PSD domains.

We initially speculated that binding of SLO to PLG would induce a large conformational change in PLG to a relaxed state to facilitate proteolytic processing by tPA. However, the HDX-MS data demonstrated that PLG does not undergo major conformational changes, at least not in a 30-9000 s temporal scale of deuteration used in this study. It should be pointed out that the disulphide bonds within PLG limits the sequence coverage in the HDX-MS analysis precluding a complete analysis of all regions of PLG. Nevertheless, HDX-MS identifies K2 and PSD domains as the primary SLO binding interfaces that overlaps with residues derived from TX-MS. In addition, we observe an exosite for SLO-binding like what has been reported for SKA or PAM interaction with PLG.^53^ The significant reduction of intra-links found within PLM compared to PLG indicates that there is a drastic conformation change upon formation of PLM that may separate the primary and secondary binding sites (exosite) in space, leading to a loss of SLO-binding as evidenced by the ELISA and the XL-MS results

The SLO-centric protein network presented here suggests novel associations with other human plasma proteins like apolipoprotein E (APOE) and clusterin (CLU). APOE contains five O-linked glycosylation sites, while CLU has six N-linked glycosylation sites. Given that cholesterol-rich eukaryotic membranes are primary targets for SLO^54,55^, it is plausible that SLO-mediated isolation of APOE occurs through interactions with cholesterol or glycans. A similar mechanism might apply to CLU, an enigmatic glycoprotein known for its broad range of glycan interactions^56,57^, which aligns with SLO ability to bind various glycans like lacto-N- neotatraose and blood group B type IV pentasaccharide on cell surfaces.^30^ Glycoproteomic reanalysis of AP-DDA data revealed significant enrichment of APOE and CLU glycopeptides in pulldown samples when using tagged SLO as bait. Recent reports indicate that human serum albumin interacts with SLO and inhibits the haemolytic effect.^32^ In our study, we did not identify albumin as a significant SLO-binder, possibly due to low affinity or competition from circulating SLO-specific antibodies. In fact, the AE-MS results show abundant enrichment of immunoglobulin peptides demonstrating that healthy plasma have numerous circulating IgG clones directed against SLO as previously shown.^58^ Previous reports have also shown that disordered GAS proteins can interact with complement components^35^. The N-terminal part of SLO is disordered, potentially explaining the specific enrichment of complement-related proteins to SLO. However, additional experimental evidence is needed to clarify if these associations are direct and indirect.

An unsolved question is whether SLO binding to the PSD domain of PLG works synergistically, cooperatively, or competitively with SKA activation *in vivo*. Further investigations are needed to clarify the nature of these interactions and its implications for GAS pathogenicity. Considering the low efficacy of tPA under normal conditions^58^, the catalysing effect exerted by SLO may hold translational therapeutic value.^59^ For example, the development of mimic peptides based on this defined binding motifs could potentially be explored as a novel approach to manipulate thrombolysis. Moreover, the discovery of this indirect PLG activation pathomechanism sheds light on the multifunctional nature of SLO. The moonlighting role proposed here augments the intimate relation between GAS pathogenicity and the host coagulation/fibrinolysis system. These findings expand our understanding of GAS virulence strategies and opens up new potential avenues for therapeutic targets against GAS infections.

## Materials and Methods

### Recombinant production of tagged-bait proteins

A composite affinity-tag, comprising hex histidine, streptavidin-binding peptide, and hemagglutinin sequences, was engineered onto the N-terminus of SLO/GFP and the C-terminus of SCPA. These tagged-GFP and tagged-SCPA proteins were synthesized, produced, and purified at the Lund University Protein Production Platform (LP3). Additionally, tagged- SLO proteins were recombinantly expressed and purified in-house. The plasmids encoding the tag-SLO fusion were synthesized and assembled by LP3, then subsequently introduced into BL21(DE3) Competent Cells (Thermo Fisher Scientific). Following o/n induction with 0.01 mM IPTG (Isopropyl ß-D-1-thiogalactopyranoside; Sigma-Aldrich) and protein extraction with BugBuster (Novagen), the target tag-SLO protein underwent Ni2+-IMAC resin (Bio-Rad) purification, following a previously established protocol^60^ alongside the manufacturer standard operating procedures. Verification of the bait proteins’ purity and sequence integrity was conducted by SDS-PAGE and bottom-up mass spectrometry (BU-MS), ensuring the successful production of the bait proteins.

### Affinity-Enrichment Mass Spectrometry (AP-MS) and data Analysis

Pooled human plasma, sourced from the Innovative Research company, was diluted with phosphate-buffered saline (PBS tablet; Sigma-Aldrich). The affinity-enrichment coupled with mass spectrometry (AP-MS) was performed in line with the workflow previously described in literature.^35^ In summary, 40 µg of the tagged-bait proteins were immobilized onto a 150 µL 50% slurry Strep-Tactin Sepharose resin (IBA Lifesciences GmbH), within a Bio-Rad spin column for each experiment. The immobilized proteins were incubated with either undiluted or 1x PBS-diluted human plasma at 37°C and 500 rpm for 2 hours, followed by rigorous washing with eight column volumes of 1x PBS buffer. Bait-purified prey proteins from plasma were then eluted with 5 mM biotin (Sigma-Aldrich) in ice-cold 1x PBS. Biotin removal was achieved by adding trichloroacetic acid (TCA; Sigma-Aldrich) to a final concentration of 25%, and samples were incubated at -20°C overnight. After centrifugation at 13,000 rpm for 30 minutes at 4°C, the supernatant was discarded, leaving the pellet, which was subsequently washed with cold acetone (Thermo Fisher Scientific). The resulting pellets were solubilized in 50 µL of 8M urea with 100 mM ammonium bicarbonate (Sigma-Aldrich), then subjected to reduction with 5 mM TCEP (tris(2-carboxyethyl)phosphine; Thermo Fisher Scientific) and alkylation with 10 mM IAA (iodoacetamide; Sigma-Aldrich), followed by an overnight digestion at 37°C and 500 rpm in a ThermoMixer (Eppendorf) using trypsin (Promega) at a 1:20 enzyme-to-substrate ratio. The digested peptides were purified using a C18 spin column (Thermo Fisher Scientific), concentrated and dried via SpeedVac (Eppendorf), and reconstituted in buffer A (2% acetonitrile, 0.2% formic acid; Thermo Fisher Scientific) for mass spectrometry analysis.

For the liquid chromatography-mass spectrometry (LC-MS) analysis, approximately 800 ng of peptides from each experiment, as determined by a NanoDrop spectrophotometer (DeNovix), were loaded onto an EASY-nLC 1200 system interfaced with a Q Exactive HF-X hybrid quadrupole-Orbitrap mass spectrometer (Thermo Fisher Scientific). Initially, peptides were concentrated using a PepMap100 C18 pre-column (75 µm x 2 cm, with 3 µm particle size; Thermo Fisher Scientific) and subsequently resolved on an EASY-Spray column (ES903; Thermo Fisher Scientific) maintained at 45°C, following the manufacturer’s instructions. The chromatographic separation was executed using two mobile phases: Solvent A with 0.1% formic acid and Solvent B comprising 0.1% formic acid in 80% acetonitrile. A linear gradient was applied, ranging from 3% to 38% of Solvent B across a 120-minute period, with a flow rate held steady at 350 nl/min. For data-dependent acquisition (DDA) strategy, it initiated with an MS1 scan covering a m/z range of 350–1650, at a resolution of 120,000 and an AGC target of 3e6, with a maximum injection time (IT) of 45 ms. This was succeeded by the top 15 MS2 scans, each with a resolution of 15,000, an AGC target of 1e5, a 30 ms IT, and a normalized collision energy (NCE) of 28. Charge states of +1, +6-8, and higher were excluded. For data-independent acquisition (DIA) approach, it mirrored the DDA LC settings but in MS analysis adopted an MS1 scan over a m/z range of 390–1210, at a resolution of 60,000, an AGC target of 3e6, and a maximum injection time of 100 ms, followed by MS2 scans within a fixed isolation window of 26.0 m/z. Here, a resolution of 30,000, an AGC target of 1e6, and an IT of 120 ms were applied, along with a NCE of 30. To assure system performance, yeast protein extract digest (Promega) standards was analysed throughout the measurement.

The initial batch of twenty-seven DDA datasets were searched against a human reference proteome database including common contaminants (UPID: UP000005640) using MaxQuant software (version 2.0.3.0) with the default settings^61^ to construct a spectral library. Based on this, nine DIA datasets were processed using the MaxDIA workflow.^62^ pGlyco 3.0^63^ was used to identify glycopeptides of PLG from AE-MS DDA runs. Label-free quantification raw data were assessed through the MiST score pipeline^37,64^ and normalized MaxLFQ intensities were processed in Perseus^65^ to discard any invalid data points, transform data, and perform statistical analysis between groups. For comprehensive data visualization, tools such as Cytoscape^66,67^, STRING network^39^, and R packages including ggplot2 and pheatmap were employed. Furthermore, the Metascape tool^68^ was utilized for performing over-representation (enrichment) analysis to explore the association among isolated proteins and to interpret the biological significance of the data.

### Direct and indirect ELISA assessment

For the quantification of specific IgG titres via ELISA, same amounts of SLO protein or SCPA were coated to MaxiSorp plates (Thermo Fisher Scientific) and incubated overnight at 4°C. Non-specific binding sites on the plates were then blocked with 1% bovine serum albumin (BSA; Sigma-Aldrich). Subsequently, the plates were added with 10 µg of either IVIG (pooled IgG isolates from population), P.IgG (a pool of IgG isolated from the plasma of a donor who recently recovered from a GAS infection), Xolair (an anti-IgE IgG; Novartis), or 1x PBS as a background reference. After a thorough washing, 100 µL of a 1:3000 dilution of HRP-conjugated protein G (Bio-Rad) and 100 µL of HRP substrate buffer (20 ml NaCitrate pH 4.5 + 1 ml ABTS + 0.4 ml H_2_O_2_; Sigma-Aldrich) were sequentially mixed with samples. The reaction developed over 30 minutes and absorbance was read at 450 nm. For indirect ELISA aimed at validating the native PLG/PLM (Thermo Fisher Scientific) specificity for SLO, equivalent quantities of PLG or PLM were first immobilized and then incubated with SLO protein, followed by a mouse monoclonal anti-SLO antibody (Abcam) and an anti-mouse HRP-conjugated secondary antibody (Bio-Rad). The procedures for substrate development and absorbance reading were the same as the direct ELISA method above.

### Plasminogen activation and plasmin activity assay

The assays to evaluate plasminogen activation and plasmin activity were conducted using a colorimetric quantification kit from Abcam. The PLG activation assay employed 96-well microtiter plates and a mimic chromogenic substrate, whose hydrolysis by plasmin at 25°C led to the release of p-nitroaniline. This distinctive final product was then quantified by measuring absorbance at 405 nm by a BMG Labtech microplate reader. Streptolysin O (SLO) was incrementally added to the samples, according to each specified concentration in four replicates. Data obtained from the assay was analysed with GraphPad Prism software (version 9.0), where initial activation velocities were derived from the linear part of the absorbance versus time-squared plot (A_405nm_/t^2^). For the plasmin activity assay, an inhibitory mix was included to negate any non-specific background that might be present in the samples. The influence of SLO on plasmin activity was examined by the addition of SLO to the reaction, following the same processes as the PLG activation assay for the substrate addition and subsequent absorbance measurement.

### SLO cytolysis inhibition assay

In assessing the inhibitory effect of PLG on SLO-mediated cytolysis, sheep red blood cells (Thermo Fisher Scientific), were initially diluted with 1x PBS. To this diluted RBC suspension, 100 ng of TCEP-reduced active SLO protein and 5 µg of PLG/PLM proteins were added. Xolair and P.IgG were also included in the assay as negative and positive control separately. The mixtures were incubated in a ThermoMixer at 37°C 300 rpm for 30 minutes. Post-incubation, the reaction plates were centrifuged, and the supernatant was transferred to fresh plates for absorbance measurement at 541 nm using a BMG Labtech microplate reader, which corresponds to the free haemoglobin concentration indicative of cell lysis. A 100% lysis rate was set using the A_541nm_ from the group of 0.6% RIPA (Thermo Fisher Scientific) lysis buffer.

### Crosslinking Mass Spectrometry (XL-MS) and data analysis

For the crosslinking experiment, in-solution crosslinking was conducted on PLG and SLO, PLM and SLO, as well as PLG and tPA combinations, each at a 1:1 molar ratio, and for a ternary complex of PLG, SLO, and tPA at ratios of 1:1:1 and 1:2:1. These proteins were mixed in 1x PBS and incubated at 37°C in a ThermoMixer with agitation at 500 rpm for one hour. Crosslinking agents, specifically DSS-H12/D12 and DSG-H6/D6 from Creative Molecules, were then individually introduced to crosslink the protein mixtures for two hours. The reactions were quenched with 4 M ammonium bicarbonate (Sigma-Aldrich), followed by conventional reduction and alkylation steps as detailed before. A two-stage enzymatic digestion with 2 hr lysyl endopeptidase (FUJIFILM Wako Chemicals U.S.A. Corporation) and o/n trypsin (Promega) was then performed to generate the crosslinked peptides.

Once prepared, the peptides were subjected to a cleaning process, desiccated, and resuspended prior to MS analysis. An Orbitrap Eclipse Tribrid Mass Spectrometer (Thermo Fisher Scientific), linked to a Nanospray source and combined with an Ultimate 3000 UPLC system (Thermo Fisher Scientific), was used for LC-MS detection for crosslinked peptides. Approximately 600 ng of peptides, quantified using a NanoDrop spectrophotometer, were loaded onto a PepMap RSLC column (Thermo Fisher Scientific) and maintained at 45°C for concentration. Duplicate injections were carried out for each sample. Column preparation and peptide loading adhered strictly to the manufacturer’s guidelines. The chromatographic separation utilized a gradient of Solvent A (0.1% formic acid) and Solvent B (0.1% formic acid in 80% acetonitrile), progressing linearly from 4% to 38% Solvent B over 90 minutes at a flow rate of 300 nl/min. The mass spectrometer operated in positive DDA mode, with an MS1 scan ranging from 400–1600 m/z at resolution of 120,000; a standard-mode AGC target and auto-mode maximum injection time, followed by a series of MS2 scans following a set cycle time of 3 s; 15,000 resolution; standard AGC target; 22 ms IT and NCE of 30. Charge states ranging from 2 to 6 were included in the analysis. LC-MS system performance was benchmarked using HeLa protein digest standards (Thermo Fisher Scientific) throughout the process.

The databases for searching crosslinked peptide encompassed PLG/PLM (UniProtID: P00747), tPA (UniProtID: P00750), and SLO sequences (UniProtID: P0DF96), in addition to common contaminants. The search parameters regarding modification related to crosslinking were configured as specified by the manufacturers. Fixed modifications were assigned to carbamidomethylated cysteines, and oxidation(M) was marked as a variable modification. Data analysis was conducted using pLink2 (version 2.3.11) and MaxLynX (version 2.0.3.0) software to extract high-confidence spectrum evidence, achieved by reducing the maximum allowed missed cleavages of peptides from 3 to 2 with the rest of parameters kept as previously described.^43,44^ The selected crosslinked peptide MS/MS spectra was then visualized using MaxLynX. DisVis^69^ analysis was performed to visualize the interaction space of tPA using all tPA-PLG-SLO heterotrimeric protein complex conformation consistent with the identified cross-links between tPA and SLO, with Cα-Cα distance set to be within 0-40 Å for comprehensive profiling.

### Hydrogen/Deuterium Exchange Mass Spectrometry (HDX-MS) and data analysis

The Hydrogen/Deuterium Exchange Mass Spectrometry (HDX-MS) setup involved the use of a LEAP H/D-X PAL™ system for automated sample preparation, connected to an LC-MS system comprising an Ultimate 3000 micro-LC linked to an Orbitrap Q Exactive Plus MS. HDX experiments were conducted on plasminogen (PLG) in both its unbound (apo) and SLO- bound (complex) states, using a 1:2 molar ratio of PLG to SLO in 10 mM PBS. The samples were labelling at various time points (0, 30, 3000, and 9000 s) at 4°C in PBS or a D_2_O-based HDX labelling buffer (Thermo Fisher Scientific). Each state from each time point repeated three times within a single continuous run. The labelling reaction was quenched by diluting the samples with 1% trifluoroacetic acid (TFA; Thermo Fisher Scientific), 0.4 M TCEP, and 4 M urea, all maintained at 4°C. Following quenching, the samples were injected for online pepsin digestion at 4°C, in a flow rate of 50 μL/min 0.1% formic acid for 4 minutes online digestion and trapping of the samples. The digested peptides were then subjected to a solid-phase extraction and washing process using 0.1% formic acid on a PepMap300 C18 trap column (Thermo Fisher Scientific), subsequently switched in-line with a reversed-phase analytical column (Hypersil GOLD). Chromatographic separation was performed at 1°C, employing a mobile phase gradient from 5% to 50% of Solvent B (95% acetonitrile/0.1% formic acid) over 8 minutes, and then from 50% to 90% over the next 5 minutes. The separated peptides were analysed on the Q Exactive Plus MS, which featured a heated electrospray ionization (HESI) source operating at a capillary temperature of 250°C. Full scan spectra were obtained with a high resolution of 70,000, using an automatic gain control target of 3e6 and a maximum ion injection time of 200 ms, covering a scan range of 300-2000 m/z. Peptide identification was achieved by analysing un-deuterated control PLG peptide samples through data-dependent MS/MS. A comprehensive peptide library, including peptide sequence, charge state, and retention time information, was established for HDX analysis using PEAKS Studio X (Bioinformatics Solutions Inc.), searching pepsin-digested, un-deuterated samples against the PLG sequence (UniProtID: P00747). HDX-MS data were then processed and analysed using HDExaminer v3.1.1 (Sierra Analytics Inc.).

Comparative analysis of PLG in SLO-bound versus unbound states was conducted using individual charge states for each identified peptide. Deuterium incorporation levels were determined based on the observed mass differences between deuterated and control peptides, without back-exchange correction using fully deuterated samples for comparison. Manual inspection of the spectra was carried out to eliminate low-scoring peptides, obvious outliers, and those with inconsistent retention times. Deuteros 2.0^70^ software was applied to perform hybrid significance tests and to demonstrate changes in deuterium uptake through kinetic uptake, barcode, redundancy, coverage, butterfly, woods, and volcano plots. This software also contributed to the projection of the coordinates of protected peptide residues onto the crystal structure of PLG, aiding in the interpretation and visualization of binding interactions and protein dynamics.

### Targeted Cross-linking Mass Spectrometry and data analysis

The computational modelling was conducted following previously established methods^71^. In summary, the AlphaFold2-Multimer^46^ was utilized for the initial prediction of the PLG-SLO structure without incorporating experimental data. The NMSim^47^ analysis was applied to assess the plausible motions of the SLO domains resulting in 5 trajectories of 500 conformers. For the prediction of SLO conformational changes, these 2,500 conformers were investigated and the best that matched the intra-protein crosslinks were selected. As for modelling PLG-SLO pairwise complex, the Rosetta^45^ software was used to generate two sets of 10,000 models based on different SLO conformational states (native form, and bend form) through the Rosetta docking protocol^45^, with the highest-scoring models being chosen for subsequent analysis. The TX-MS^34^ technique was then employed to select top-ranking models based on distance constraints derived from XL-MS datasets. Finally, a high-resolution refinement of the top-ranking models was performed using the RosettaDock protocol^72^ to repack sidechain. The binding interfaces were then defined using the PRODIGY tool.^45^

## Data Availability

All mass spectrometry proteomics data have been deposited to the ProteomeXchange Consortium via the PRIDE^73^ partner repository with the dataset identifier PXD051261 (The dataset will be made publicly available upon completion of the peer review process for this manuscript.).

## Conflict of Interest Disclosure

The author has no competing interests to declare.

## Acknowledgements

We gratefully acknowledge the Swedish National Infrastructure for Biological Mass Spectrometry (BioMS), the SciLifeLab Integrated Structural Biology Platform, and the Protein Production Sweden (PPS) for providing facilities and experimental support. We would also like to thank Dr. Hong Yan, Dr. Tommaso De Marchi, Dr. Alejandro Gomez Toledo, Dr. Annika Rogstam and Dr. Wolfgang Knecht for assistance.

## Funding

J.M. is a Wallenberg academy fellow (KAW 2017.0271) and is also funded by the Swedish Research Council (Vetenskapsrådet, VR) (2019-01646 and 2018-05795), the Wallenberg foundation (KAW 2016.0023, KAW 2019.0353 and KAW 2020.0299), and Alfred Österlunds Foundation. H.K. is supported by the French Agence Nationale de la Recherche (ANR), under grant ANR-22-CPJ2-0075-01.

---

**Extended Data Figure 1.**
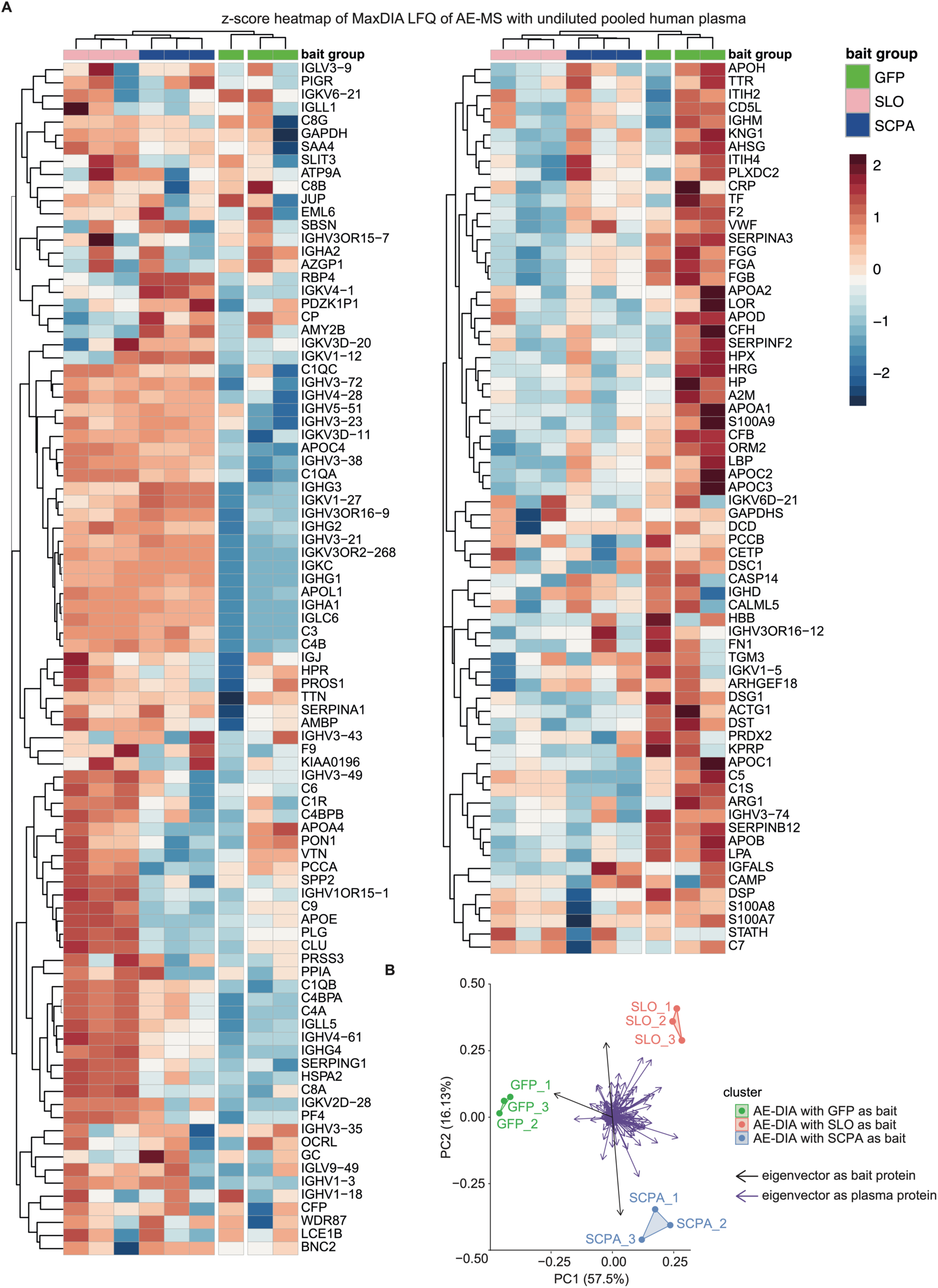
Proteomics profiling of all identified human plasma proteins through AE-MS DIA. **A)** Heatmap of z-score normalized AE-DIA data. Intensities of all proteins from three bait groups (SLO, SCPA and GFP) across three replicates were analysed, with the colour scale indicating protein group enrichment (red) or reduction (blue) in comparison across conditions. **B)** PCA plot clusters nine samples based on protein abundances, with bait protein contribution marked in black arrow.

**Extended Data Figure 2.**
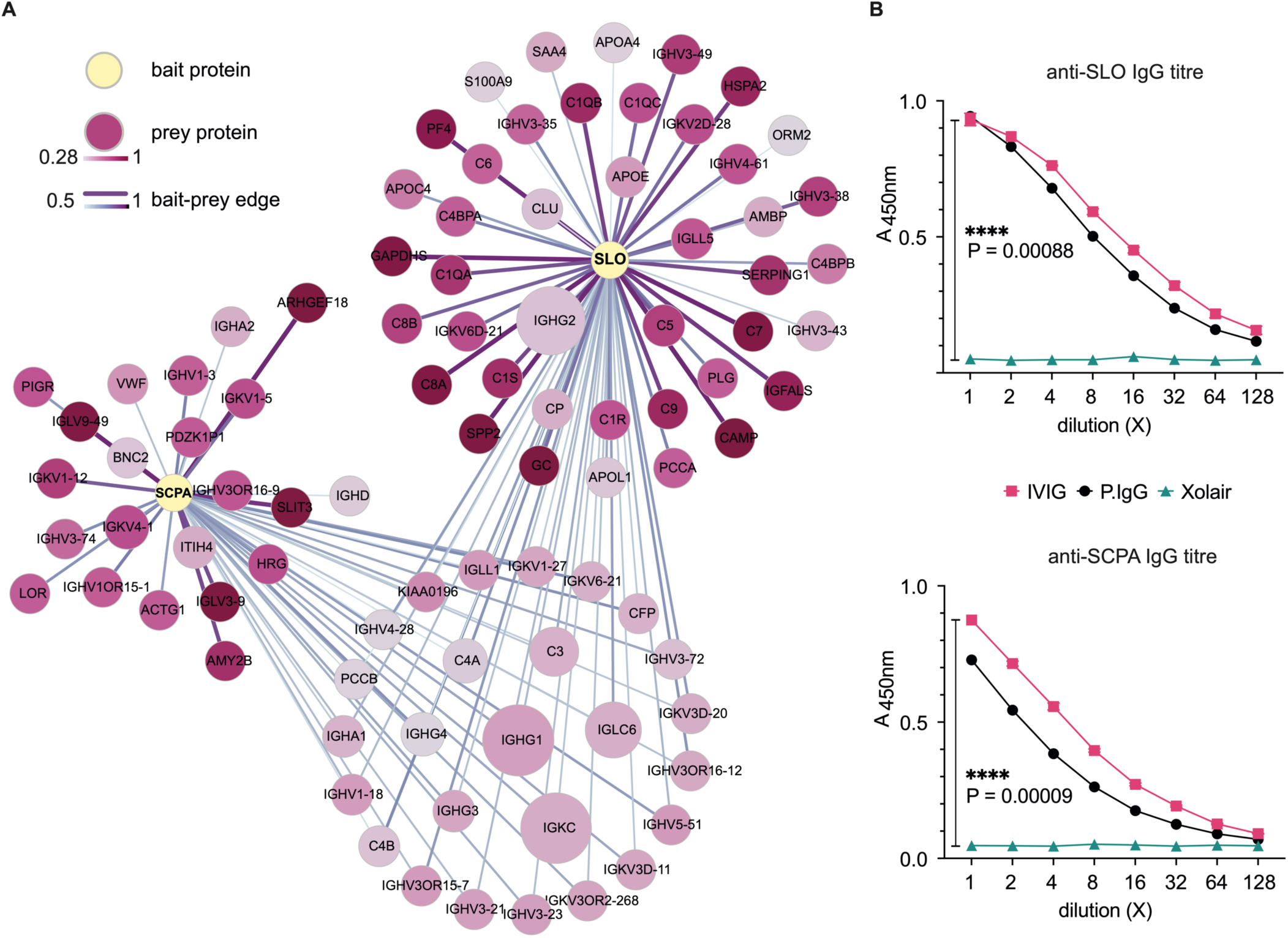
Combined PPI networks of SLO/SCPA and plasma protein, and circulating specific IgGs in plasma. **A)** This network depicts the interactions between tagged-SLO/tagged-SCPA and human plasma proteins. Nodes represent proteins, with size and colour indicating abundance and specificity of the isolated prey protein, while connecting edges, representing protein-protein interactions (PPI), are defined by width and colour based on calculated MiST scores. **B)** ELISA measurement of IgG titres against SLO (streptolysin O) and SCPA (C5a peptidase) in different IgG samples. Absorbance values at 450 nm (A_450nm_) are shown for IVIG (pooled human IgG isolates), P.IgG (IgGs from a patient recovering from a recent GAS infection), and Xolair (negative control, a monoclonal IgG1 specific to IgE).

**Extended Data Figure 3.**
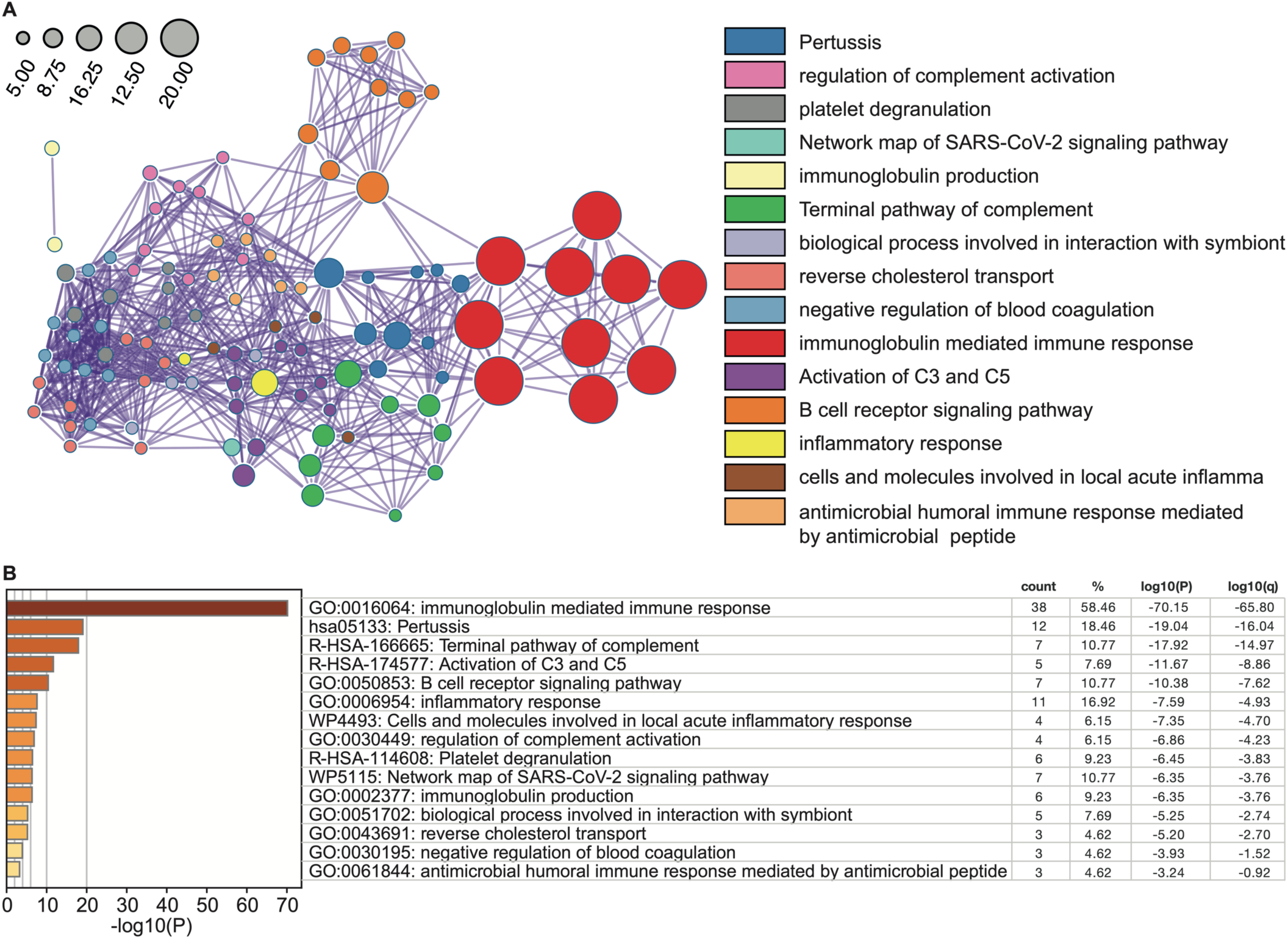
Over-representation analysis of SLO-enriched proteins. Based on MiST scores, 68 putative SLO-interactive plasma proteins are applied here for over-representation analysis. **A)** The network visualization represents enriched terms from the gene ontology biological processes (GO-BP), KEGG pathway, Reactome, and WikiPathways databases. The most term from the top 15 enriched clusters is displayed. **B)** The bar graph displays these 15 representative enriched terms, with colours indicating the p-values, coupled with the “Count” as the number of genes from the input list associated with the given term, the “%” representing the proportion of input genes in the term, and the log-transformed p-values “Log10(P)” and multi- test adjusted p-values “Log10(q)” for statistical context.

**Extended Data Figure 4.**
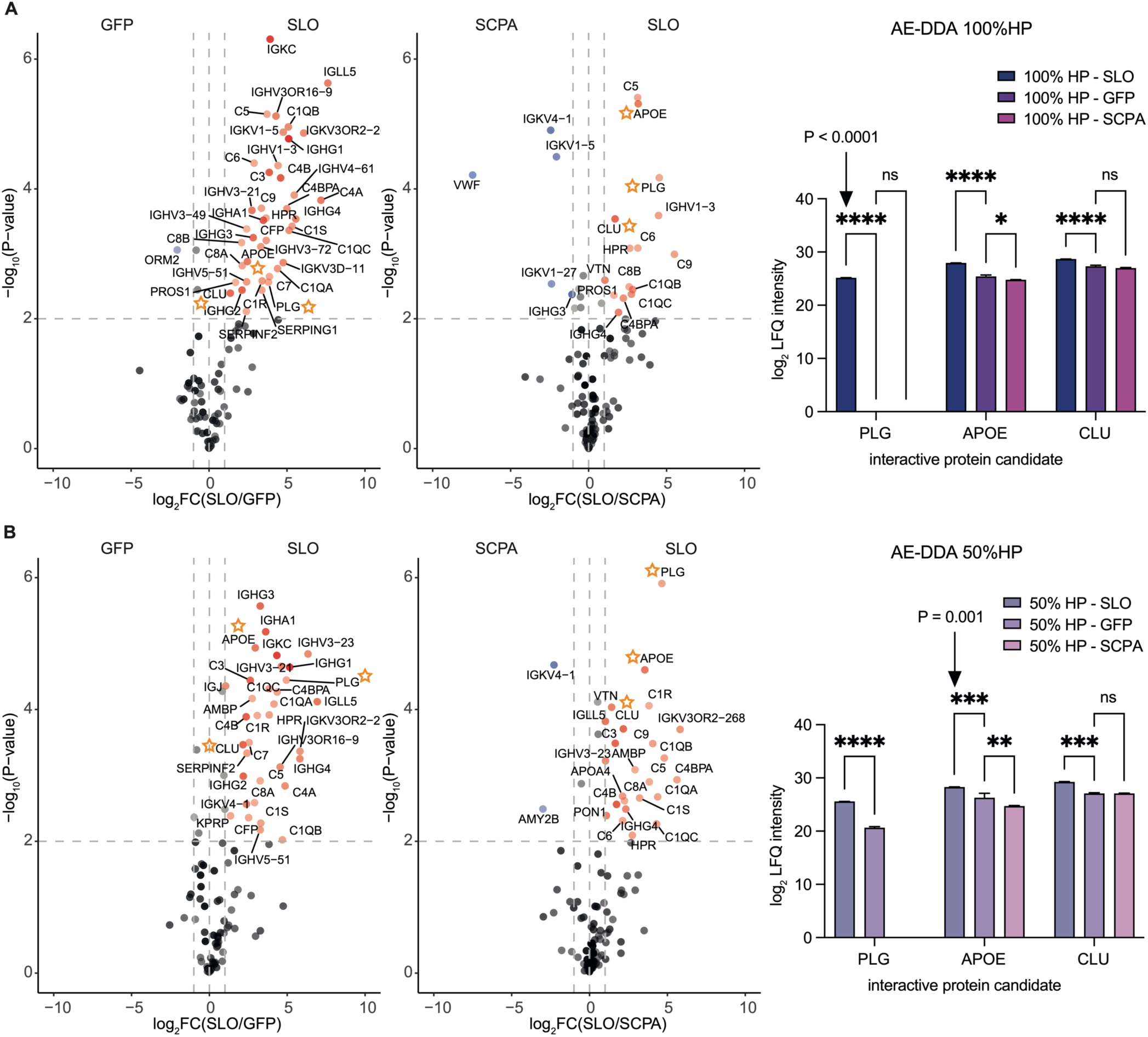
AE-DDA MS differential analysis using diluted human plasma as prey mixture. Pooled human plasma, prepared undiluted (100%) or diluted to 50% with PBS, was subjected to AE-MS experiment in three independent replicates. Proteins eluted were analysed via DDA and searched against the reference proteome database. Prey protein abundances in SLO bait group were compared against GFP or SCPA group. Fold change of normalized LFQ data and multiple t-test statistics identified proteins differentially enriched by SLO, coloured in red. Volcano plots were plotted for **A,** left) undiluted and **B,** left) 50% diluted group accordingly. Two-way ANOVA on log2-transformed MaxLFQ intensities highlighted the top three putative SLO-interactive plasma proteins: plasminogen (PLG), apolipoprotein E (APOE), and clusterin (CLU). **A-B,** right) Histograms display the mean and standard deviation of candidate protein intensities, with varying colour depths indicating dilution factors. Significance levels are denoted as “ns” (not significant), “*”, “**”, “***”, and “****” for P-values < 0.1, < 0.01, < 0.001, and < 0.0001, respectively.

**Extended Data Figure 5.**
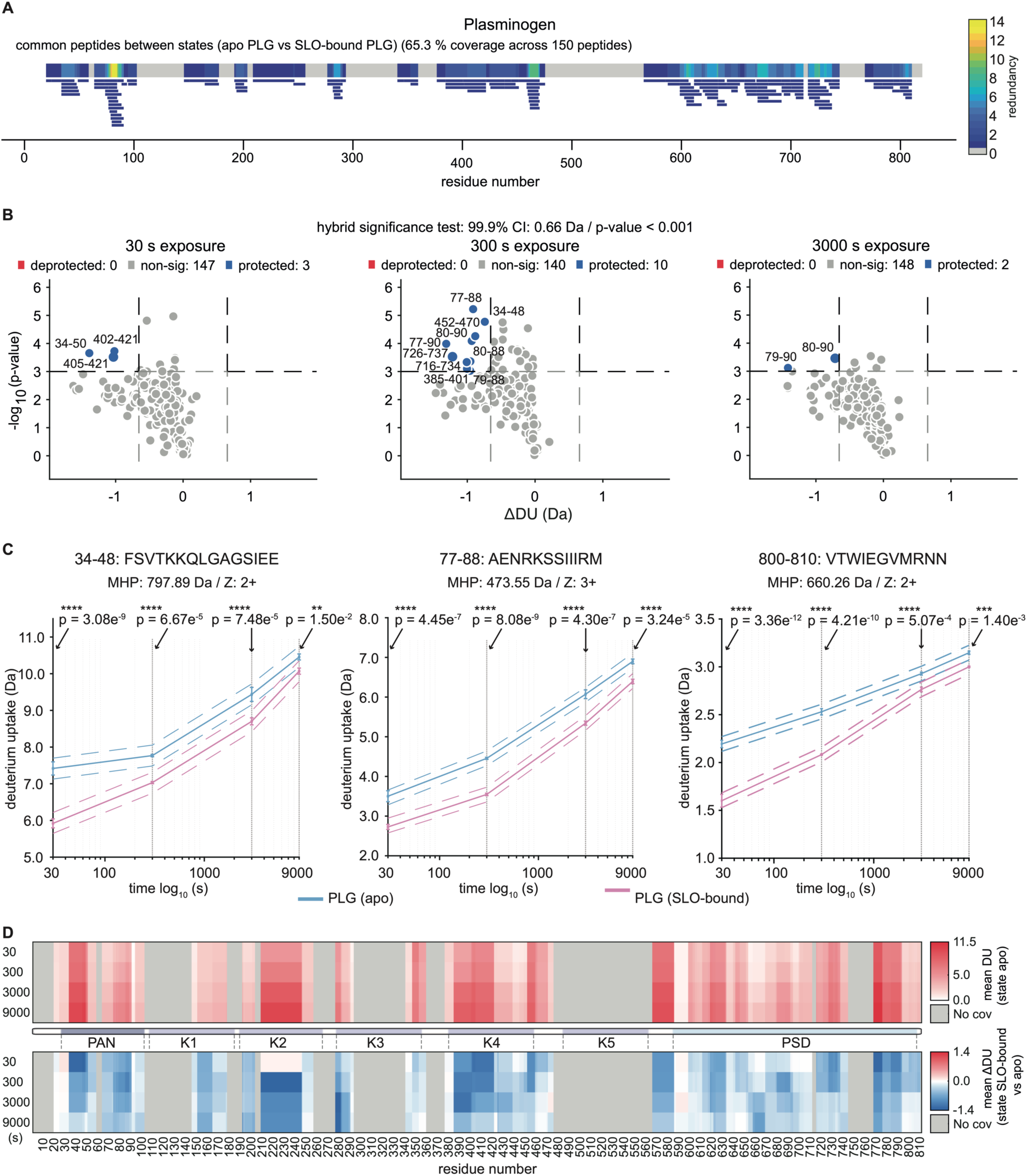
Protected regions and protein dynamics of SLO-bound PLG revealed by HDX-MS. **A)** A redundancy map for 150 identified peptides common to both states of apo PLG and the SLO-bound PLG in HDX-MS, with bars representing peptides and the sequence gradient indicating residue redundancy. **B)** Volcano plots display peptides with significant changes in deuterium uptake at 30, 300, and 3000 s, and peptides are plotted as dots, coloured based on its protection/deprotection status. **C)** Kinetic plots for three protected peptides in PAN and PSD region, with significance testing across four labelling times, color-coded to different states. Significance levels are marked, with p-values annotated. ’MHP’ refers to theoretical molecular weight of the peptide, “Z” to charge state. **D)** The top panel barcode plot display the mean deuterium uptake of PLG residues in apo state while the bottom panel represent mean differential deuterium uptake in SLO-bound state, with the colour gradient indicating the change level, as a function of labelling time.

**Extended Data Figure 6.**
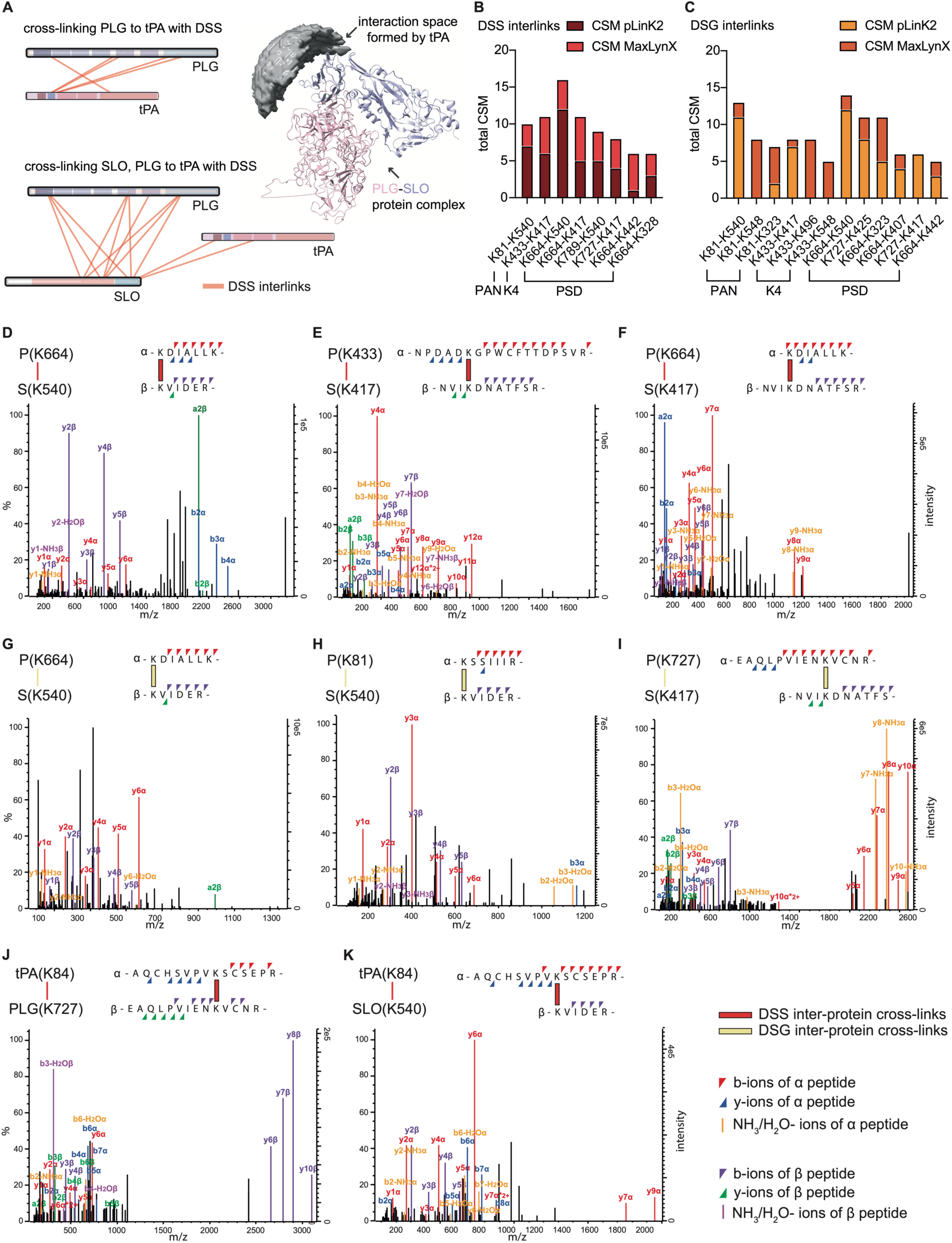
Linkage maps of inter-links for PLG-tPA and PLG-SLO-tPA, CSM count summary of inter-links and annotated MS/MS spectra of cross-linked peptide pairs found in PLG-SLO, PLG-tPA and tPA-SLO Interactions. CSM counts between PLG and SLO from **A)** Two linkage maps display the DSS inter-protein cross-links, revealing interactions within the PLG-tPA and PLG-SLO-tPA complexes. The interaction space around PLG-SLO complex was formed by all tPA-PLG-SLO conformations consistent with two reported cross-link distance constraints between tPA and SLO. **B)** DSS cross-linked dataset and **C)** DSG cross-linked dataset, from both search engines. **D-I)** present the MS/MS spectra of the representative peptide pairs between PLG domains (PAN, Kringle 4, PSD) and SLO. **J)** showcases one MS/MS spectra of cross-linked peptide pairs between tPA and PLG. **K)** displays one MS/MS spectra of cross-linked peptide pairs between tPA and SLO. All fragmented ions are annotated and coloured for clarity.

**Extended Data Figure 7.**
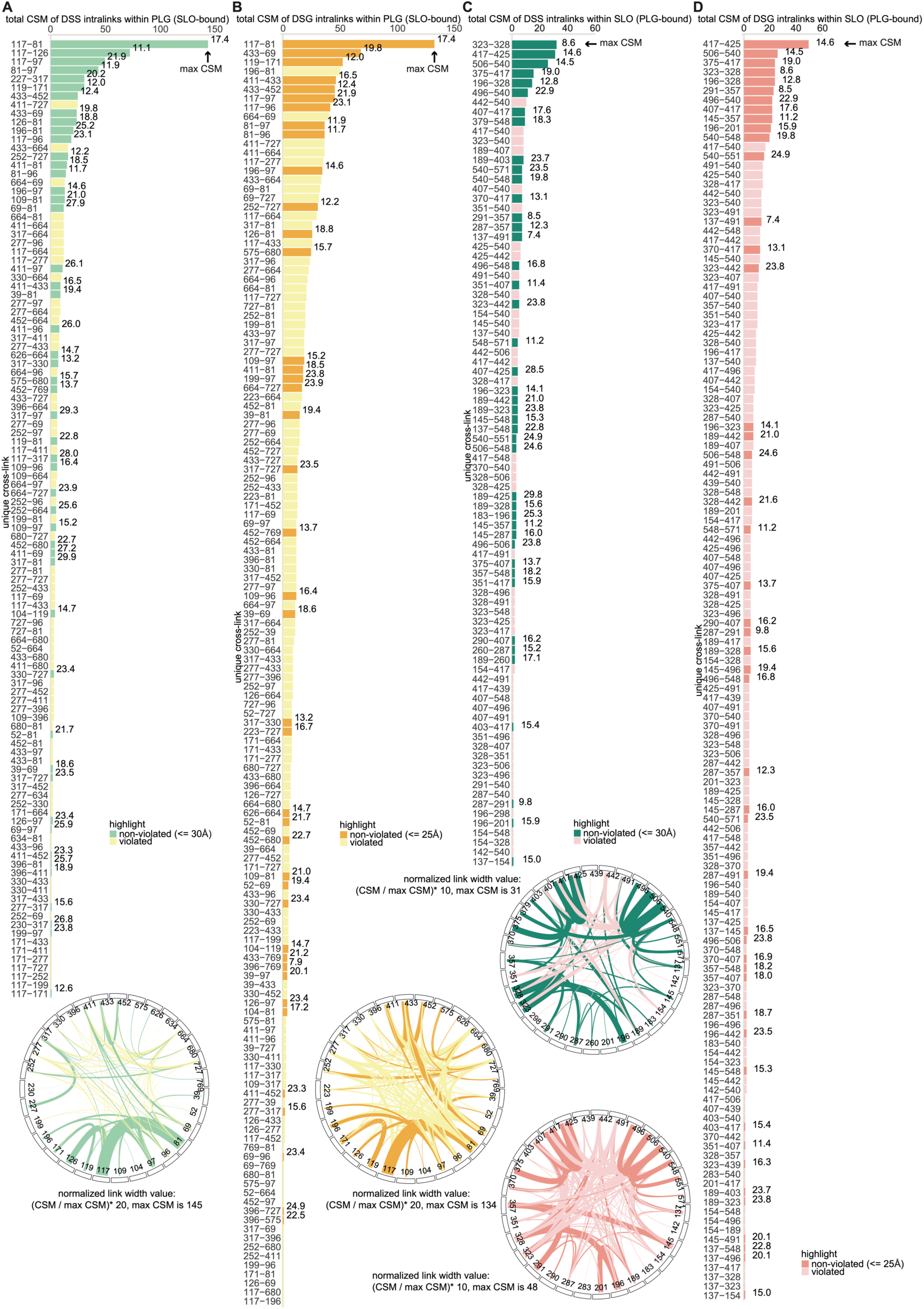
CSM summary of intra-links found in PLG or SLO during interaction. **A-D)** Bar plots present CSM count across all unique XLs, and circos plots depict XL sites with edges width indicating CSM counts.

**Extended Data Figure 8.**
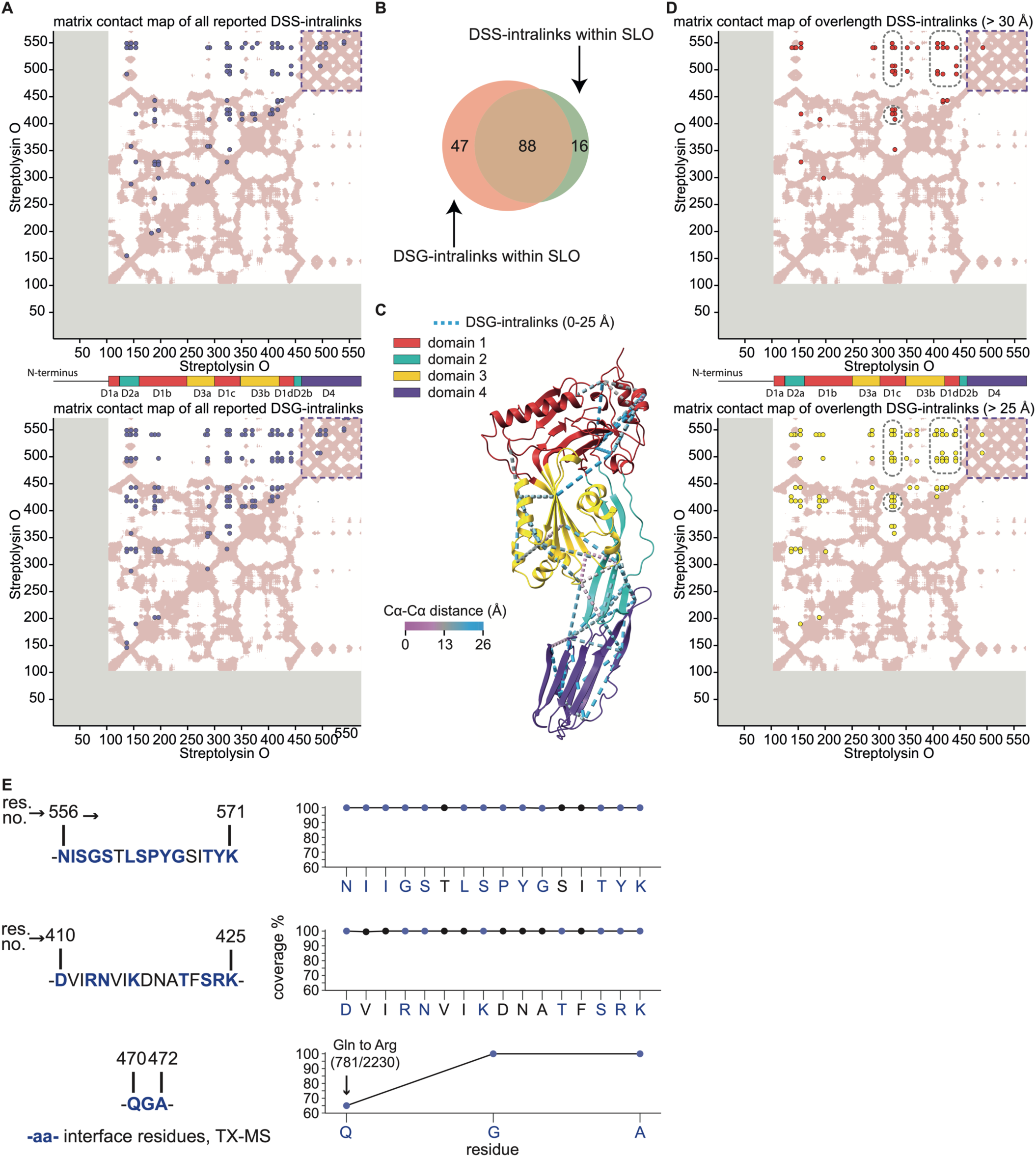
Matrix contact map of intra-links found within SLO against its reference crystal structure. **A)** Two matrix map presents all identified intra-protein cross-links in dots against a context of SLO structure (PDB: 4HSC). White background represents sequence coverage in input structure, grey for absence and pink for ambiguous cross-link or match (Cα-Cα proximity < 25 Å). Structured regions of SLO, 103-571, are coloured by domain separately, with a-d representing subsequence. **B)** A Venn diagram details the overlap and unique intra-protein cross-linked sites found within SLO protein from both DSS and DSG cross-linked datasets. **C)** A crystal structure of SLO is coloured by domain, with consistent DSG intra-links (distance < 25 Å) displayed in dot-line style pseudo bond. **D)** Two matrix map presents all only distance-violating intraprotein cross-links (>30 Å for DSS; >25 Å for DSG) in dots against a context of SLO structure. **E)** Three PLG-binding interfaces on SLO protein were derived from TX-MS top-ranked model, and analysed for residues conservation among 2230 high-quality GAS genomes sourced from The Bacterial and Viral Bioinformatics Resource Centre. Streptolysin O sequence (Uniprot ID: P0DF96) is set as reference. Interface contact residues derived from TX-MS are coloured in blue and marked in bold.

**Supplementary Data Figure 1.**
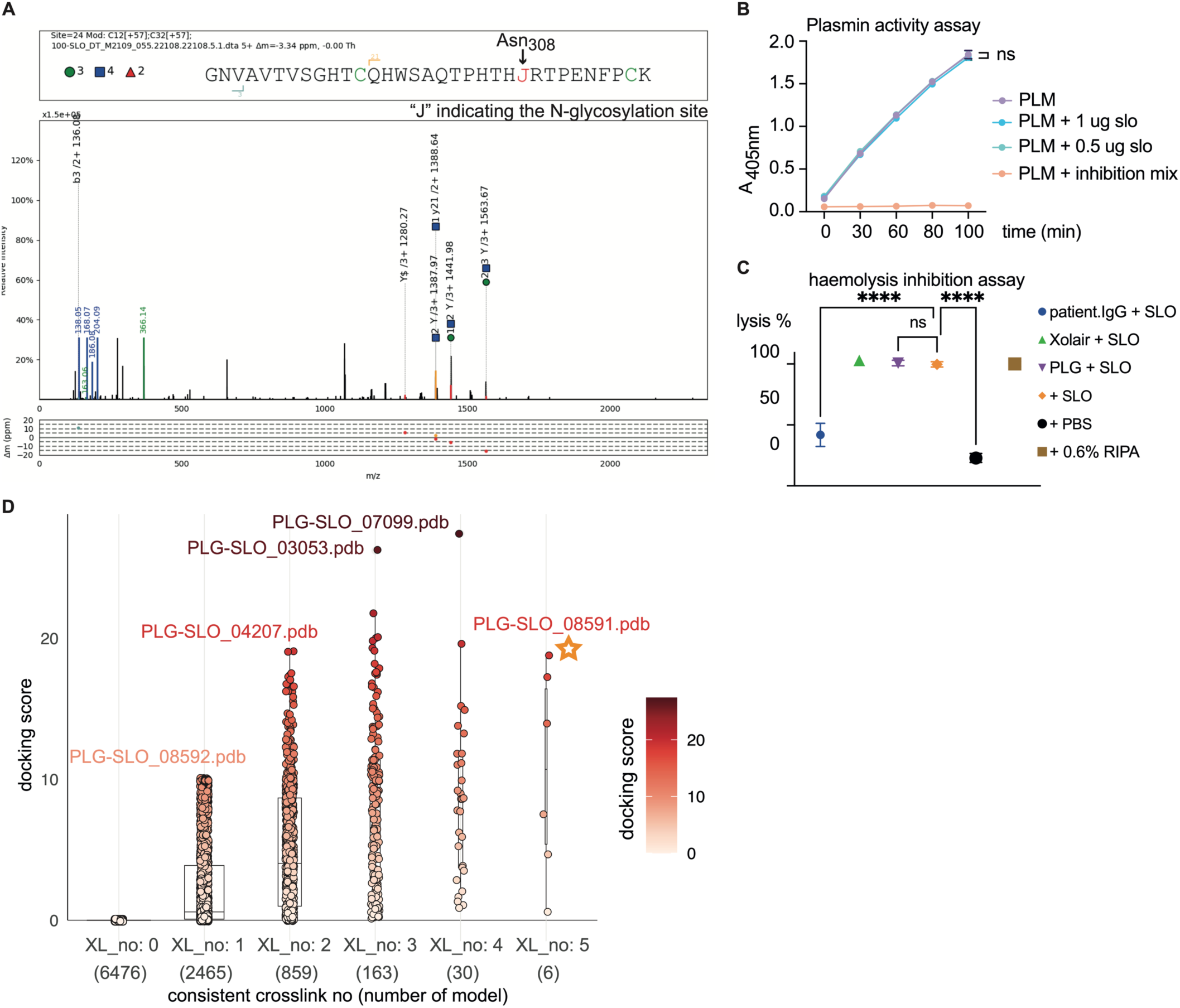
Identification of PLG glycopeptides in SLO-enriched samples and related functional assays. **A)** Here displays a MS/MS spectra of one representative PLG glycopeptide identified in SLO-enriched plasma samples. The analysis, performed with pGlyco 3.0 and visualized via pGlycoLabel, revealed glycosylation at Asn308 on PLG, indicative of type 1 PLG. **B)** Plasmin activity assay was performed where SLO and plasmin (PLM) were incubated for 30 minutes before adding a chromogenic plasmin substrate. The graph plots absorbance at 405 nm against the time of development, including three replicates per condition. **C)** SLO-mediated haemolysis assay was conducted where SLO, pre-incubated with various proteins, was activated by TCEP and then added to the diluted suspension of sheep red blood cells. RIPA, a cell lysis detergent, acted as the positive control. Haemolysis percentage was determined based on haemoglobin levels of supernatant after centrifuge in the experimental groups relative to the positive control. Statistical differences among groups were evaluated using one-way ANOVA, with significance levels marked as “ns” (not significant), “*”,“**”, “***”, and “****” for P-values < 0.1, < 0.01, < 0.001, and < 0.0001, respectively. **D)** The box plot and accompanying individual dot plot depict the distribution of docking scores across all pairwise PLG-SLO(bend form) models. The number of models that satisfy the inter-protein cross-links is annotated along the x-axis.

**Supplementary Data Table 1.** CSM Summary of inter-protein cross-links between PLG and SLO. The table lists all searched DSS and DSG inter-protein cross-linked sites between P (Plasminogen) and S (streptolysin O), ranked separately by the total number of CSM reported by pLink2 (rank_pL) and MaxLynX (rank_Ma) search engines, with a CSM sum (SUM CSM) and an aggregated ranking order (SUM rank).

| P-S; DSS<br>inter-links | CSM_<br>pLink2 | rank_<br>pL | CSM_<br>MaxLynx | rank_<br>Ma | SUM<br>CSM | SUM<br>rank |  | P-S; DSG<br>inter-links | CSM_<br>pLink2 | rank_<br>pL | CSM_<br>MaxLynx | rank_<br>Ma | SUM<br>CSM | SUM<br>rank |
| --- | --- | --- | --- | --- | --- | --- | --- | --- | --- | --- | --- | --- | --- | --- |
| K664-K540 | 12 | 1 | 4 | 3 | 16 | 4 |  | K664-K540 | 12 | 1 | 2 | 5 | 14 | 6 |
| K664-K417 | 5 | 4 | 6 | 1 | 11 | 5 |  | K81-K540 | 11 | 2 | 2 | 5 | 13 | 7 |
| K433-K417 | 6 | 3 | 5 | 2 | 11 | 5 |  | K727-K425 | 8 | 3 | 3 | 4 | 11 | 7 |
| K81-K540 | 7 | 2 | 3 | 4 | 10 | 6 |  | K664-K323 | 5 | 6 | 6 | 2 | 11 | 8 |
| K789-K540 | 5 | 4 | 4 | 3 | 9 | 7 |  | K433-K417 | 7 | 4 | 1 | 6 | 8 | 10 |
| K727-K417 | 4 | 5 | 4 | 3 | 8 | 8 |  | K81-K548 |  | 11 | 8 | 1 | 8 | 12 |
| K664-K442 | 1 | 8 | 5 | 2 | 6 | 10 |  | K433-K496 |  | 11 | 8 | 1 | 8 | 12 |
| K664-K328 | 3 | 6 | 3 | 4 | 6 | 10 |  | K81-K323 | 2 | 9 | 5 | 3 | 7 | 12 |
| K664-K491 | 2 | 7 | 2 | 5 | 4 | 12 |  | K664-K407 | 4 | 7 | 2 | 5 | 6 | 12 |
| K81-K496 |  | 9 | 3 | 4 | 3 | 13 |  | K727-K417 | 6 | 5 |  | 7 | 6 | 12 |
| K277-K540 |  | 9 | 3 | 4 | 3 | 13 |  | K664-K442 | 3 | 8 | 2 | 5 | 5 | 13 |
| K396-K540 |  | 9 | 3 | 4 | 3 | 13 |  | K433-K548 |  | 11 | 5 | 3 | 5 | 14 |
| K433-K328 | 2 | 7 | 1 | 6 | 3 | 13 |  | K81-K417 | 1 | 10 | 3 | 4 | 4 | 14 |
| K433-K323 |  | 9 | 2 | 5 | 2 | 14 |  | K81-K491 | 1 | 10 | 3 | 4 | 4 | 14 |
| K727-K323 |  | 9 | 2 | 5 | 2 | 14 |  | K433-K425 | 1 | 10 | 3 | 4 | 4 | 14 |
| K727-K540 |  | 9 | 2 | 5 | 2 | 14 |  | K727-K328 | 1 | 10 | 3 | 4 | 4 | 14 |
| K789-K496 |  | 9 | 2 | 5 | 2 | 14 |  | K81-K442 | 2 | 9 | 2 | 5 | 4 | 14 |
| K727-K442 | 1 | 8 | 1 | 6 | 2 | 14 |  | K81-K407 | 3 | 8 | 1 | 6 | 4 | 14 |
| K664-K496 | 1 | 8 | 1 | 6 | 2 | 14 |  | K171-K189 | 1 | 10 | 2 | 5 | 3 | 15 |
| K97-K540 | 1 | 8 | 1 | 6 | 2 | 14 |  | K433-K540 | 1 | 10 | 2 | 5 | 3 | 15 |
| K96-K540 | 1 | 8 | 1 | 6 | 2 | 14 |  | K433-K491 | 2 | 9 | 1 | 6 | 3 | 15 |
| K81-K442 | 1 | 8 | 1 | 6 | 2 | 14 |  | K727-K442 | 2 | 9 | 1 | 6 | 3 | 15 |
| K81-K328 |  | 9 | 1 | 6 | 1 | 15 |  | K433-K323 | 3 | 8 |  | 7 | 3 | 15 |
| K81-K506 |  | 9 | 1 | 6 | 1 | 15 |  | K664-K548 |  | 11 | 2 | 5 | 2 | 16 |
| K81-K548 |  | 9 | 1 | 6 | 1 | 15 |  | K664-K417 | 1 | 10 | 1 | 6 | 2 | 16 |
| K97-K298 |  | 9 | 1 | 6 | 1 | 15 |  | K96-K496 |  | 11 | 1 | 6 | 1 | 17 |
| K117-K540 |  | 9 | 1 | 6 | 1 | 15 |  | K664-K496 | 1 | 10 |  | 7 | 1 | 17 |
| K171-K496 |  | 9 | 1 | 6 | 1 | 15 |  | K330-K417 |  | 11 |  | 7 | 0 | 18 |
| K330-K417 |  | 9 | 1 | 6 | 1 | 15 |  |  |  |  |  |  |  |  |
| K397-K407 |  | 9 | 1 | 6 | 1 | 15 |  |  |  |  |  |  |  |  |
| K397-K442 |  | 9 | 1 | 6 | 1 | 15 |  |  |  |  |  |  |  |  |

**Supplementary Data Table 2.**
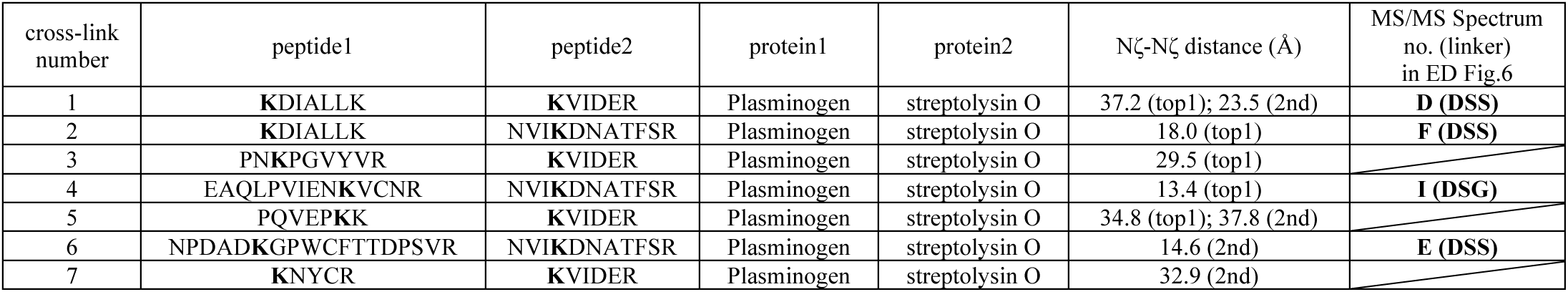
Summary of inter-protein cross-links between PLG and SLO. The table presents the identified cross-linked peptide pairs between PLG and SLO, specifying the numbering of cross-links. For each XL peptide pair (Peptide 1 and Peptide 2), the reactive lysine residues are highlighted in bold. The table also denotes the source proteins (Protein 1 and Protein 2) for the respective peptides. The distance between the cross-linked lysine side chains (Nζ-Nζ) is provided for both top1 model shown in Figure 5C and 2nd model in Figure 5E, with a reference to the corresponding MS/MS spectra, which can be found in **Extended Data Figure 6** if available.

## Notes

### Competing Interest Statement

The authors have declared no competing interest.

